# Disentangling single-cell omics representation with a power spectral density-based feature extraction

**DOI:** 10.1101/2021.10.25.465657

**Authors:** Seid Miad Zandavi, Forrest Koch, Abhishek Vijayan, Fabio Zanini, Fa Valdes Mora, David Gallego Ortega, Fatemeh Vafaee

## Abstract

Emerging single-cell technologies provide high-resolution measurements of distinct cellular modalities opening new avenues for generating detailed cellular atlases of many and diverse tissues. The high dimensionality, sparsity, and inaccuracy of single cell sequencing measurements, however, can obscure discriminatory information, mask cellular subtype variations and complicate downstream analyses which can limit our understanding of cell function and tissue heterogeneity. Here, we present a novel pre-processing method (scPSD) inspired by *power spectral density* analysis that enhances the accuracy for cell subtype separation from large-scale single-cell omics data. We comprehensively benchmarked our method on a wide range of single-cell RNA-sequencing datasets and showed that scPSD pre-processing, while being fast and scalable, significantly reduces data complexity, enhances cell-type separation, and enables rare cell identification. Additionally, we applied scPSD to transcriptomics and chromatin accessibility cell atlases and demonstrated its capacity to discriminate over 100 cell types across the whole organism and across different modalities of single-cell omics data.

## Main

Continuous innovations in single-cell technologies allow the interrogation of a growing number of molecular modalities such as DNA, chromatin, mRNA and protein, at high-resolution and across thousands of cells from complex biological systems. Increased throughput of new single-cell technologies has posed unique analytical challenges demanding for scalable computational methods that can analyze diverse high-dimensional omics data highly accurately and fast^1^. Single-cell sequencing data also suffer from the ‘curse of missingness’ due to, for instance, dropout events in scRNA-sequencing^2^ or the low copy number in DNA leading to an inherent per-cell sparsity in scATAC-sequencing data^3^. High-dimensionality and sparsity, combined with various systematic biases in single-cell sequencing experiments^4^, obscure important information in data which hinders precise distinctions among cell states and masks shared biological signals among different cell subtypes. Extracting discriminatory information is therefore essential for the success and accuracy of downstream analyses and is particularly relevant for the application of machine learning methods to diverse problems from cell-type classification to trajectory inference or multimodal data integration^5^.

Feature extraction seeks an optimal transformation of the input data into a latent feature vector with the primary goal of extracting important information from input data, controlling for confounding effects, adjusting over-dispersion, and removing redundancy to enhance the separation of distinct cellular phenotypes^6^. Dimensionality reduction (DR) techniques such as PCA (principal component analysis)^7^, t-SNE (t-distributed stochastic neighbor embedding)^8^, and UMAP (Uniform Manifold Approximation and Projection)^9^ are frequently employed to transform high-dimensional data into a low-dimensional space, which is particularly useful to visually inspect the distribution of input data. Further feature extraction methods were specifically developed for scRNA-sequencing data – e.g., ZIFA (zero inflated factor analysis)^10^, ZinbWave (zero-inflated negative binomial model)^11^, and scVI (single-cell variational inference)^12^ – or to a much lesser extent, for other single-cell modalities – e.g., SCALE (single-cell ATAC-seq analysis via latent feature extraction)^13^. DR methods have varied performance in separating biological clusters as per our recent comprehensive benchmarking^14^ and often perform poorly in facilitating the detection of rare cell populations^15^. Furthermore, the capacity of different DR methods in extracting features from other single-cell omics, beyond scRNA-sequencing data, is undetermined and yet to be assessed systematically.

Here, we present an innovative unified strategy for single-cell omics data transformation (scPSD) that is inspired by *power spectral density* (PSD) analysis^16^ to intensify discriminatory information from single-cell genomic features. PSD is a statistical signal processing technique to describe the distribution of power over frequency and to show the strength of the energy as a function of frequency^16^. One purpose of estimating spectral density is to detect any patterns or periodicities in a signal by observing peaks at the frequencies corresponding to these patterns. Here, a vector of genomic features (e.g., expressions of transcripts, open chromatin regions, or cell-surface proteins in a single cell) has been realized as a ‘signal’ representing a cellular state. The scPSD feature transformation performs four consecutive steps on ‘single-cell genomic signals’ (**Fig 1a**):

**Fig 1.**
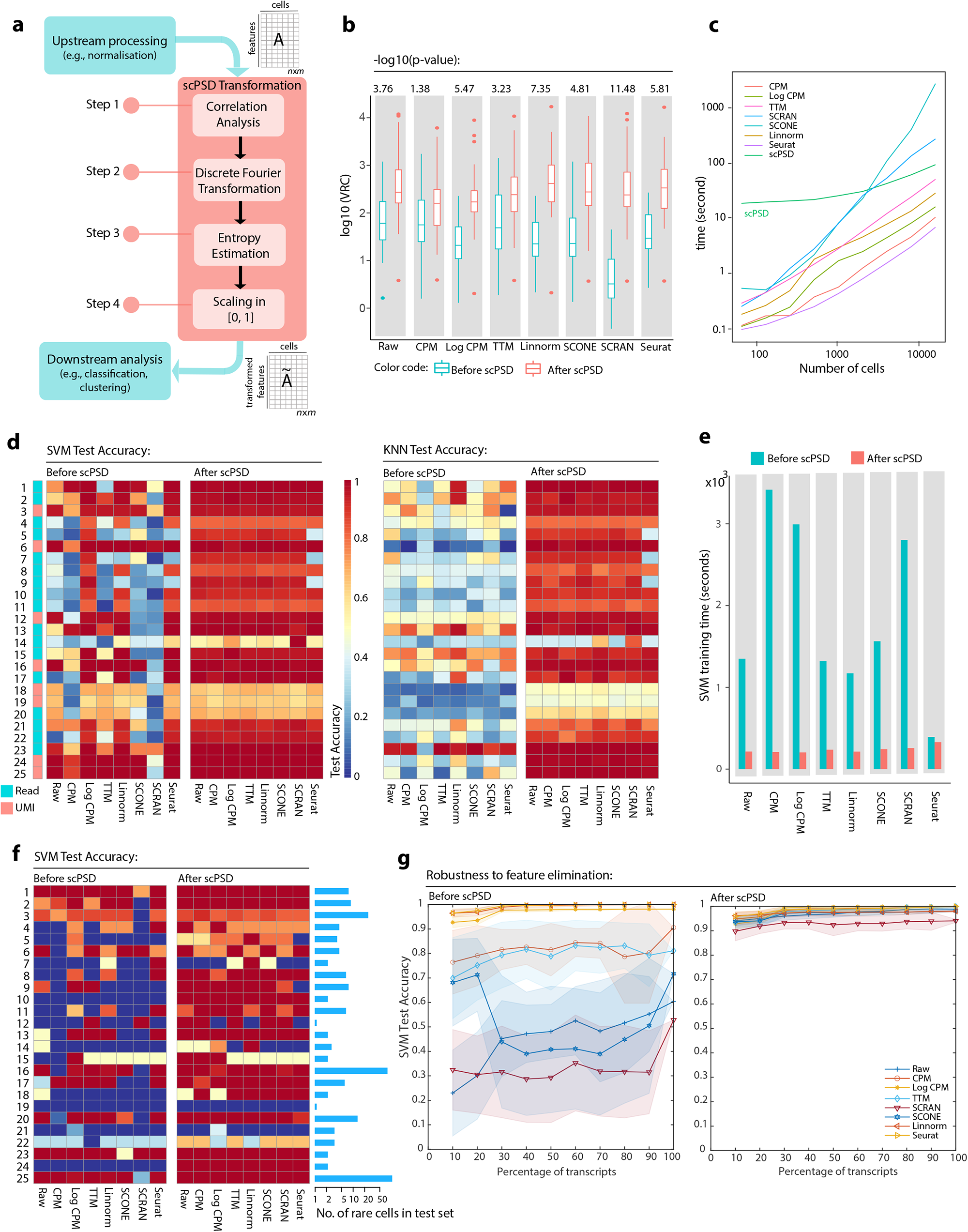
Overview of scPSD and performance evaluation on scRNA-seq datasets. **a**, the scPSD transformation framework (red rectangle) comprising four consecutive steps of feature extraction and standardization. scPSD can fit into a single-cell sequencing analysis pipeline after the upstream processing (or directly on raw data) to enhance downstream analyses. **b**, box plots comparing VRC (variance ratio criterion) as a measure of how well-formed distinct cell-types are before/after scPSD transformation of normalized and raw counts across 25 curated scRNA-seq datasets (numbered according to Supplementary Table 1); p-values of t-tests comparing log10-transformed VRC measures before and after scPSD are reported on top of each pair of boxplots. SS (silhouette score) and mFDR (multi-class Fisher’s discriminant ratio) as other measures of cluster separation and dataset complexity are reported in Supplementary Fig 2. **c**, computational runtime of scPSD and normalization methods as scales with increasing number of cells. Methods were applied to datasets of varying size obtained by random subsampling of the 10X Genomics E18 mouse dataset, and timings are averaged over 16 applications. **d**, heatmaps representing accuracy of cell-type prediction – for each of 25 scRNA-seq datasets – on 20% randomly held out data (test set) after training SVM (support vector machine) and KNN (k-nearest neighbor) models on remaining 80% of data (training set), before and after scPSD transformation. **e**, SVM training time in second before and after scPSD transformation demonstrating significant reduction in convergence time after transformation. **f**, heatmaps representing SVM test accuracy identifying rare cell-type identification – defined as the smallest cell-type population constituting <1% to 14% of captured cells across 25 scRNA-seq datasets – before and after scPSD transformation. **g**, SVM test accuracy upon increasing feature coverage using ‘deng reads’ dataset (#13 in Supplementary Table 1). The procedure includes random accumulation of genes (in 10% brackets), reporting SVM test accuracy (on 20% holdout cells) before and after scPSD transformation, and repeating the procedure 100 times to account for random feature selection. The average trends were reported with shades representing +/-standard deviation across 100 repeats.

**1)** Estimating pairwise correlations of genomic features across cells followed by within-cell correlation mapping. **2)** Feature extraction by discrete Fourier transformation (DFT), a mathematical approach widely used to reveal hidden patterns and periodicities across a finite data sequence upon transformation into the frequency domain. As reviewed elsewhere^6^, DFT has been used in a variety of bioinformatics applications for the analysis of repetitive elements in DNA sequences and protein structures, among others. We implemented the fast Fourier transform (FFT), a highly efficient procedure for computing the DFT of a data sequence^17^. **3)** Entropy estimation to improve the extraction of important information from Fourier transformed data. We employed Shannon’s entropy^18^ which describes the uncertainty in discrete random variables representing the information content of a probabilistic event. Entropy-based methods have been frequently used for feature extraction and analysis of biological sequences as reviewed previously^19,20^. **4)** Scaling transformed values between zero and one.

The scPSD transformation can fit into any single-cell computational pipeline complementing the upstream pre-processing (e.g., normalization) to improve data quality, and streamlining downstream computations (**Fig 1a**) and is independent of any initial random ordering of genomic features (**Supplementary Fig 1**).

We assessed the performance of scPSD transformation, independent of any downstream analyses, in improving the cell-type clustering tendency and reducing the complexity of single-cell omics data. We previously proposed^21^ a supervised application of internal validation measures (IVMs) such as silhouette score (SS)^22^ and variance ratio criterion (VRC)^23^, to quantify the compactness and separation of annotated cell-type clusters. We also defined a measure of the complexity of a multi-class dataset inspired by the Fisher’s discriminant ratio (FDR)^24^ (detailed in online Methods) to quantify the pairwise difference and dispersion of individual features among different cell types.

We comprehensively evaluated the effect of scPSD transformation in improving clustering tendency and complexity of single-cell transcriptomics data across 25 scRNA-seq datasets representing 14 tissue types, 10 sequencing protocols that resolved between 4 and 56 distinct cell-types (**Supplementary Table 1**). Upon selecting these datasets, we carefully assessed the underlying cell-type determination approaches (detailed in **Supplementary Table 1**) to incorporate trustworthy annotations for a reliable performance evaluation.

We have shown that scPSD significantly improves data quality (as measured by SS, VRC and our modified FDR) over these 25 RNA-seq datasets (**Fig 1b** and **Supplementary Fig 2-3**) while being efficient in time (**Fig 1c**). We applied scPSD on raw and normalized data. Multiple commonly-used normalization methods such as trimmed means of M-values (TMM)^25^, count per million (CPM)^26^, and seurat^27^ as well as single-cell-specific methods namely scone^28^, Linnorm^29^, and scran^30^ were used as a pre-processing step prior to the transformation. We observed that while the type of normalization significantly affects data quality before transformation, after conducting scPSD, the quality of transformed data is not influenced by the normalization method, i.e., *p*-value of ANOVA test across different normalization methods on log-transformed VRC measures is significant before scPSD transformation (*p* = 3.57E-11), but insignificant afterwards (*p* = 0.478). This shows that scPSD transformation not only reduces the complexity of datasets for downstream analyses, but also can be used to harmonize single-cell omics data derived from diverse preprocessing pipelines for reuse, integration, and construction of cell atlases.

Feature extraction is essential to improve the performance of machine learning algorithms. Supervised classification methods, for instance, have been widely adopted for automatic cell labelling to predict the identity of each cell by learning from an annotated training data^31^. We compared the performance of the general-purpose support vector machine (SVM), the best performing classifier based on a former benchmarking study on scRNA-seq data^31^, as well as other commonly used classifiers (i.e., random forest (RF) and k-nearest neighbor (KNN)) before and after scPSD feature extraction. For each dataset, the performance was evaluated based on the classification accuracy over a holdout test set (20% random split of a dataset) as well as the training computation time. The results clearly support a significant improvement in both metrics (due to the reduced complexity and faster convergence) irrespective of the choice of classifier or the upstream pre-processing approach (**Fig 1d-e**, and **Supplementary Fig 4**).

We further examined the effectiveness of scPSD transformation in facilitating the detection of rare cell populations. Rare or low abundant cell types within complex tissues can play important roles in normal development or disease progression (e.g., stem and progenitor cells, and circulating tumour cells)^32^. Therefore, identifying rare cell populations can be of significant interest and the performance of rare cell type identification may not be consistent with the general classification performance. We reported the percentage of correctly classified cells belonging to the smallest cell-type population in each dataset, ranging from 2 – 500 cells, before and after scPSD transformation on raw and normalized datasets. We observed a clear improvement in identifying rare cells after transformation (**Fig 1f** and confusion matrices in **Supplementary Fig 5**).

We originally trained classifiers on the full set of genes. Classifiers, however, are often sensitive to the number of features (genes) used^31^ necessitating a careful feature selection prior to classification. To assess the sensitivity of the classification performance to the number of features, we randomly selected 10% of genes from a modest-sized scRNA-seq dataset (deng reads^33^) and obtained SVM test accuracy before and after scPSD transformation. We then added another 10% of holdout genes, obtained the accuracy, and continued until accumulating all genes. This whole procedure was repeated 100 times to account for the random nature of the feature selection. Interestingly, we observed that the classifier has become extremely robust to feature elimination/selection upon scPSD transformation (**Fig 1g**).

Furthermore, to compare the effectiveness of scPSD feature extraction with feature extraction via dimensionality reduction, we studied 33 DR methods (**Supplementary Table 4**), extracted low-dimensional latent features across multiple datasets, and assessed the clustering tendency of cell types as measured by VRC and SS. We observed significantly higher IVMs using scPSD transformed features compared to features obtained by different DR methods (**Supplementary Fig 6**). Overall, scPSD can be used as a standalone feature extraction or precede a DR method to enable visualization (**Supplementary Fig 3**) or reduce data for more efficient downstream analyses.

Beyond cell-type classification and clustering tendency, we assessed the utility of scPSD in improving developmental trajectory inference (TI). We studied 20 datasets representing diverse trajectory types, i.e., linear, bifurcation, multifurcation, and tree (**Supplementary Table 5**) and used minimum spanning tree (MST), a previously-shown^34^ well-performing method, to infer topologies. As recommended by Saelens *et al*^34^, we used multiple metrics for comparing trajectories including the Hamming–Ipsen– Mikhailov (HIM)^35^ metric, F1 between branch assignments, and correlation between geodesic distances (detailed in Methods). Our initial results (**Supplementary Figure 7**) show that the MST average performance across all datasets measured by different metrics significantly improved after applying scPSD transformation (paired t-test p-value < 0.01) though the method performance was variable across datasets with different trajectory types (**Supplementary Table 6**).

In addition, we assessed scPSD performance on three in-house, well-characterized single cell transcriptomics datasets (**Supplementary Table 2**) where scRNA-seq combined with fluorescent multiplexed in situ hybridization and flow cytometry were used to characterize changes in composition of immune cells^36^ (5,234 cells and 16 cell subtypes), mesenchymal cells^37^ (5,479 cells and 16 subtypes), and endothelial cells^38^ (2,930 cells and 10 subtypes) in the murine lung during early postnatal development. Across three datasets, sequencing reads were obtained following the same protocol (Smart-Seq2) and bioinformatics pipeline resulting identical sequencing coverage. We applied scPSD on the CPM-normalized combined dataset (including 13,643 cells, 42 subtypes, and 18,072 transcripts) and observed improvement in separation and dispersion of distinct cell-types (measured by SS and VRC) after scPSD feature extraction (**Fig 2a**). Strikingly, scPSD transformation disentangled the representation of the complex landscape of proliferative macrophages comprising multiple types of macrophages, dendritic cells, granulocytes, and lymphocytes as visualized by t-SNE 2D scatter plots before and after scPSD transformation (**Fig 2a**).

**Fig 2.**
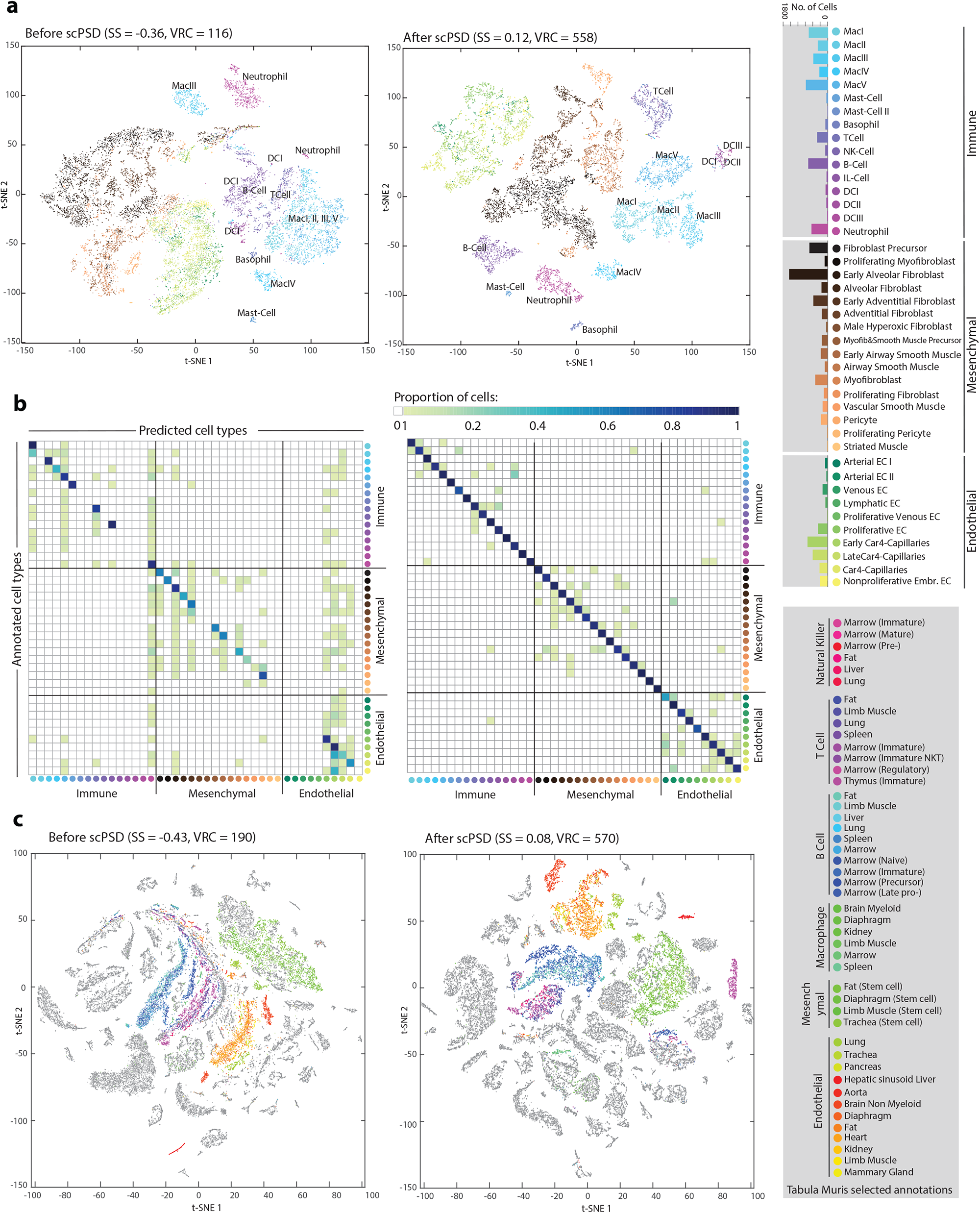
Evaluating scPSD performance on an atlas of the murine lung immune compartment and Tabula Muris mouse transcriptomic cell atlas. **a**, t-SNE 2D visualizations (plus SS and VRC measures) before and after scPSD transformation of in-house scRNA-seq profiles capturing murine lung immune cell landscape combined with mesenchymal and endothelial cell subpopulations during postnatal development (data detailed in Supplementary Table 2). The combined profile was CPM normalized prior to visualization and transformation; immune cell subpopulations were annotated on the plots. **b**, confusion matrices detailing the performance of SVM classification on test set (20% randomly held out cells) before and after scPSD transformation. Rows represent annotations (i.e., true classes) while columns represent predictions. Confusion matrices report the proportion of false positives, false negatives, true positives, and true negatives allowing more detailed analysis of cell-specific miss-classification. **c**, t-SNE visualizations of Tabula Muris transcriptomics cell atlas accompanied with clustering tendency metrics (SS and VRC) for qualitative and quantitative evaluation of scPSD transformation on an entire organism. The immune, mesenchymal, and endothelial cells from different tissues were color-coded, other cells were grayed out however are explorable via interactive Supplementary Files 1 and 2.

Confusion matrices in **Fig 2b** detail the cell-specific (mis-)classification rate on test set (20% holdout samples) using an SVM classifier trained on 80% of the combined dataset. Interestingly, while scPSD significantly improves classification performance (**Supplementary Table 3**), misclassified cells are often within the same cellular category in contrast to the pre-scPSD prediction where, for instance, multiple immune or mesenchymal cells were predicted as endothelial cells. Of note, since scPSD is an unsupervised procedure (i.e., does not rely on cell annotations), misclassified cells may also indicate occasional errors in the original annotations.

Beyond individual organs, we corroborated the efficacy of scPSD to process the single-cell transcriptomic data of an entire organism, the *Tabula Muris* mouse cell atlas^39^, comprising over 100,000 cells from 20 organs and tissues. The scPSD transformation was efficient (84 seconds using 16 CPUs on UNSW HPC platform, Katana) and enhanced cell-type separation as measured by VRC and SS (**Fig 2c**). Interestingly, the t-SNE 2D plots show close proximities, yet often with distinct boundaries, in latent feature space among cell types that are shared between tissues, e.g., immune, mesenchymal, and endothelial cells from different anatomical locations (**Fig 2c**). To enable further visual investigation of relationships between cells from different organs, the interactive t-SNE plots of *Tabula Muris*, before and after scPSD transformation, were made available as **Supplementary Files 1-2**.

As another level of validation, we used a CITE-seq dataset^40^ of bone marrow cells measuring transcriptomes in parallel with 25 cell-surface proteins representing well-characterized markers. The protein expression clearly discriminates immune subpopulations (**Figure 3a**) and can be considered as a gold standard for enumerating cell subsets based on quantitative differences in surface markers. While RNA-sequencing measures cannot differentiate cell-subtypes *a priori*, scPSD transformation enables high-resolution separation of cell types in concert with marker-based immunophenotyping (**Figure 3a**).

**Fig 3.**
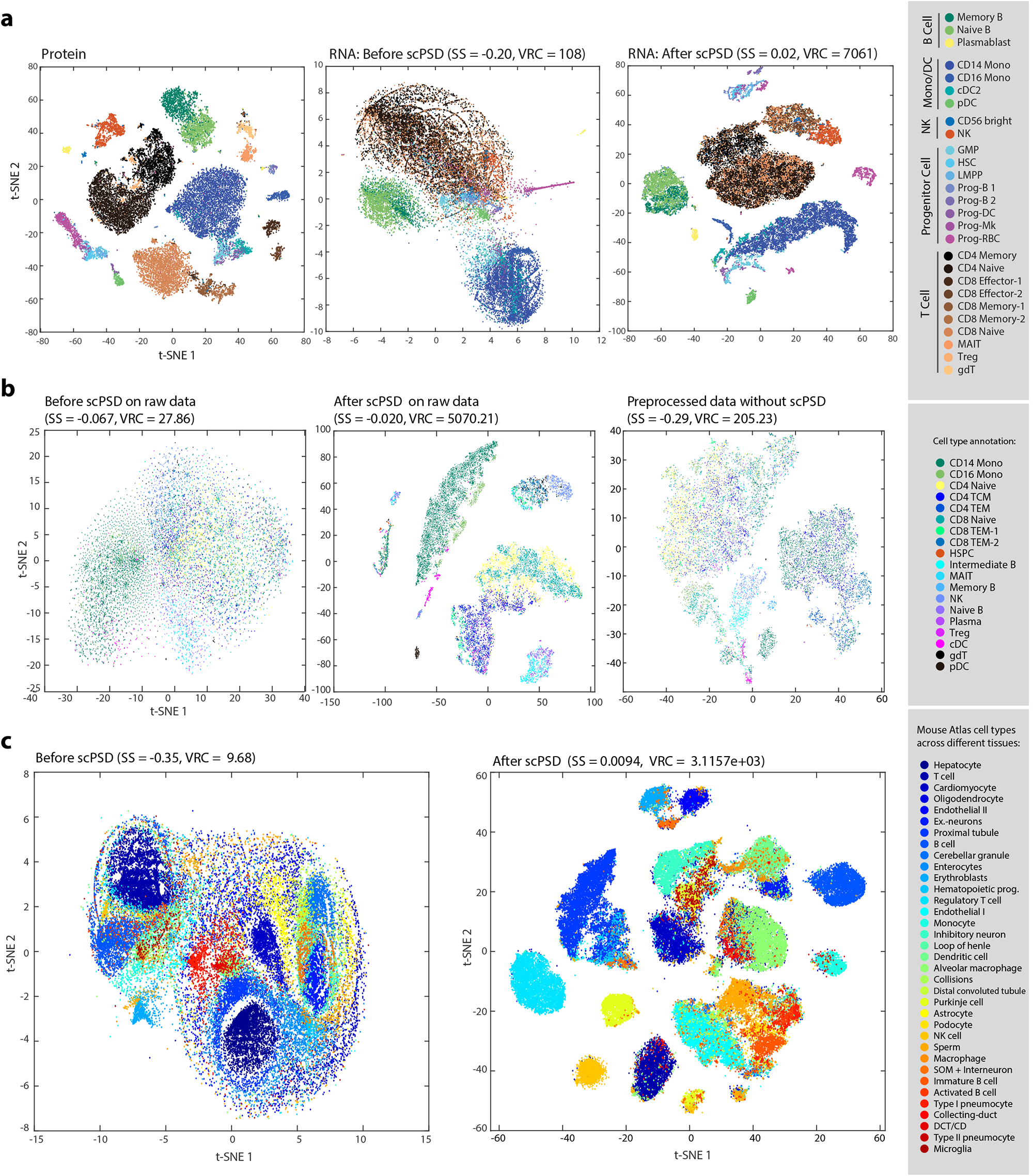
Performance validation on a CITE-seq dataset; evaluating scPSD applicability to scATAC-seq datasets. **a**. t-SNE 2D plots of a CITE-seq dataset^40^ of bone marrow cells where immunophenotypes are measured in parallel with transcriptomes including the visualization of cell subpopulations based on cell-surface protein measurements (left plot), CPM-normalized scRNA-seq profiles before scPSD transformation (middle plot) and afterwards (right plot). **b**. t-SNE visualizations of a 10X Genomics scATAC-seq dataset of human PBMC (downloaded from ‘[filtered feature barcode matrix](https://support.10xgenomics.com/single-cell-multiome-atac-gex/datasets/1.0.0/pbmc_granulocyte_sorted_10k) (HDF5))’ without pre-processing via normalization and feature selection (left plot), after scPSD transformation on non-processed data (middle plot) and after preprocessing including normalization and feature selection using ‘RunTFIDF’ and ‘FindTopFeatures’ (with q75 cutoff) functions implemented by signac’ library in R (right plot) **c**. t-SNE visualizations of mouse single-cell atlas of chromatin accessibility before and after scPSD transformation; for both plots profiles are TFIDF normalized and features with zero values in more than 95% of cells were filtered out prior to visualization and transformation. The corresponding interactive plots are available as Supplementary Files 3-4.

Beyond single cell transcriptomics, the scPSD transformation is expected to be applicable to other single-cell omics modalities as well as bulk sequencing data. As a proof of principle, we applied scPSD on two scATAC-seq datasets to assess its performance on single-cell measurements of chromatin accessibility. First, we analyzed the 10x Genomics scATAC-seq dataset of human PBMC granulocytes comprising 108,377 peaks across 11,909 cells. **Fig 3b** shows the t-SNE visualizations of the raw profile, preprocessed data (TFIDF normalization followed by inclusion of top 25% most common features using *signac* library in R^41^), and raw data after scPSD transformation. Together, the results demonstrate that although pre-processing enhances the separation of cell subpopulations, scPSD transformation further improves cell subtype clustering both visually and quantitatively (as measured by SS and VRC, **Fig 3b**). We further applied scPSD on the single-cell atlas of chromatin accessibility in mouse capturing ∼100,000 cells across 13 different tissues and similarly observed that the scPSD transformation significantly improves cell-type separation quantitatively (measured by SS and VRC) and qualitatively (shown by t-SNE plots), c.f., **Fig 3c**, and **Supplementary Files 3-4** for interactive plots.

## Methods

### Overview of scPSD

A signal can be considered as a series of measurements that conveys information about the behavior of a system. Inspired by the idea that a cell is a biological system whose behavior can be realized by a collective quantification of pools of molecules (i.e., omics), we considered an ‘omics signal’ of length *n*, denoted *a* =(*a*_1_, *a*_2_,…, *a*_*n*_) ∈ ℝ^*n*^, as a series of molecular measurements (ordered in any random arrangement).

Any signal can be decomposed into a number of discrete frequencies according to Fourier analysis. The statistical average of the signal in terms of its frequency content is called its spectrum which often contains essential information about the nature of the signal and behavior of the system. The power spectral density (PSD), or simply power spectrum, describes the distribution of *energy* into frequency components composing a signal where energy is defined as the area under the squared magnitude of the considered signal^16^. Power spectral density, therefore, indicates energetic frequencies to extract patterns and periodicities of signal. The power spectral density can be found as the Fourier transform of the autocorrelation function^42^. Autocorrelation is the correlation of a signal with a delayed copy of itself as a function of delay and depends on the ordering of datapoints in a series. Since molecular measurements (omics) are not necessarily ordered, we relaxed the ordering dependency by estimating between and within-sample correlations followed by discrete Fourier transformation. These two steps were followed by entropy estimation and scaling to get final transformed features. Accordingly, scPSD implements the following four consecutive steps (after filtering genes with zero expression across all cells):

#### Step 1. Correlation estimation

Let’s denote a single cell omics dataset as a matrix *A*=(*a*_*kj*_) ∈ ℝ^*n*×*m*^ of *n* measurements (e.g., genes) and *m* samples (i.e., cells). Across-sample correlation is obtained by computing pairwise linear correlation coefficient between each pair of genes: *ρ* = corr(*A*^*T*^) =(*ρ*_*kj*_) ∈ ℝ^*n*×*n*^ such that

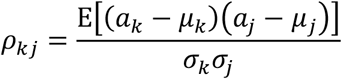

where *μ*_*k*_, *σ*_*k*_ and *μ*_*j*_, *σ*_*j*_ are means and standard deviations for genes *k* and *j* across samples. Each sample (i.e., a column vector of *n* measurements) is then linearly transformed^43^ by the correlation matrix resembling within-sample autocorrelation implemented by a matrix multiplication, *A*_1_ = *ρ* × *A*∈ ℝ^*n*×*m*^.

#### Step 2. Discrete Fourier transformation

Each column of *A*_1_, representing transformed molecular measurements of a cell, is then undergone discrete *Fourier transformation (DFT)*. We used the fast Fourier transform (FFT)^17^, an efficient method for computing the DFT. For vectors *X* and *Y* of length *n*, DFT transformation, *Y* = *DFT*(*X*) is defined as^44^

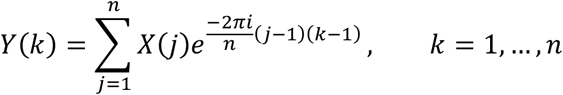

where *e*^2*πi*/*n*^ is a primitive *n*^*th*^ root of 1. Accordingly, *A*_2_ = |DFT(|*A*_1_|)|/*n* ∈ ℝ^*n*×*m*^ would be the absolute value of the fast Fourier transformation of *A*_1_ representing the ‘power’ per unit of signal.

#### *Step 3. Entropy estimati*on

As another level of feature extraction, the entropy of each sample’s probability distribution was estimated. The probability distribution of sample/cell *j* was simply estimated via scaling each feature by the sample’s marginal sum, 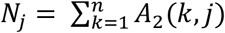, resulting probability matrix *P* ∈ ℝ^*n*×*m*^ where *P*(*k,j*) = *A*_2_(*k, j*)/*N*_*j*_. Accordingly, the entropy of sample *j* is calculated as

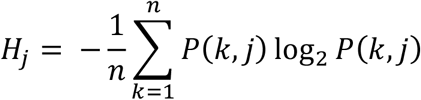

Finally, *A*_3_ ∈ ℝ^*n*×*m*^ is estimated as *A*_3_(*k, j*) = *H*_*j*_ + *P*(*k, j*) log_2_ *P*(*k, j*) for *k* = 1,…,*n* and *j* = 1,…,*m*.

#### Step 4. Scaling between zero and one

Finally, the values of each sample represented in columns of *A*_3_ will be scaled so that its range is in the interval [0,1].

### Internal Validation Measures

To quantify the compactness and separation of annotated cell-type clusters prior to any downstream analysis, we calculated internal validation measures (IVMs) for groups of cells defined by cell-type annotations provided with each published dataset. Two measures were used for this purpose: silhouette score (SS) and variance ratio criterion (VRC) as defined below:

*Variance Ratio Criterion*: VRC^21^ is the ratio of between-cluster dispersion to within-cluster dispersion and is defined as per equation below where *k* is the number of clusters, *n* is the number of data points, BGSS is the between group sum-of-squares, and WGSS is the within group sum-of-squares. Larger values of VRC indicate high dispersion between clusters and low dispersion within clusters.

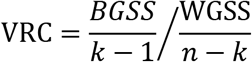

#### Silhouette Score

SS is calculated as the mean silhouette coefficient over the dataset, and varies between −1 and 1 with larger values being better^20^. A high silhouette score indicates that each point is more similar to points in its own cluster than to points from other clusters. Assume that data have been clustered via any technique (or as per annotations) into *k* clusters (i.e., cell types). For each point *i* in cluster *C*_*i*_ (i.e., *i* ∈*C*_*i*_ assuming |*C*_*i*_| > 1), the silhouette coefficient is defined as:

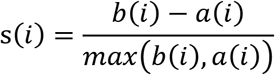

where *a*(*i*) is the average distance, *d*(*i, j*), of point *i* to each other point within the same cluster, *C*_*i*_, and *b*(*i*) is the average nearest-neighbor distance to each cluster formulated as:

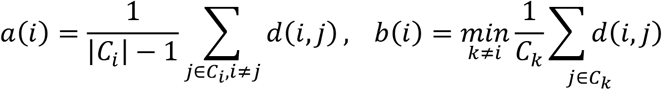

### Multiclass Fisher’s Discriminant Ratio

The Fisher’s discriminant ratio, *f*_*ij*_,was used as a separability measure of two classes of *i* and *j* and defined as^24^

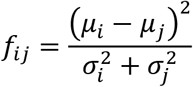

where *μ*_*i*_, 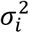 and *μ*_*j*_, 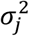 are means and variances for classes *i* and *j*. Consider *f*_*i*_ = (*f*_*i*1_, *f*_*i*2_, …, *f*_*iN*_) a vector representing the pairwise Fisher’s discriminant ratio between class *i* and *j* for *j* = 1 … *N*, where *N* is the total number of classes (cell types). We defined *F*_*i*_ as the approximate integral of *f*_*i*_ estimated via the trapezoidal integration implemented by *trapz* function in MATLAB or R. Finally, the multi-class Fisher’s discrimination ratio (mFDR) was calculated as

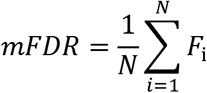

### Normalization

As a pre-processing step prior to the scPSD transformation, multiple commonly used bulk RNA-seq normalization methods as well as single-cell-specific methods were used including:

*Trimmed means of M-values* (*TMM*) ^25^ which estimates the scaling factor based on the overall expression fold-change between the sample and a reference sample. The reference sample is the one which has an upper quartile closest to the mean upper quartile of all samples. TMM is implemented in the Bioconductor R package *edgeR*.

*Count Per Million* (*CPM*) ^26^ uses as the scaling factor the sum of the read counts across all transcripts in a sample multiplied by one million.

*Scone*^28^ assesses the efficacy of various normalization workflows prior to finalizing their data normalization strategy. Scone is implemented in the Bioconductor R package *scone*^28^. The default setting was chosen to select a top ranked method among scone library wrapper^28^, upper-quartile scaling normalization^45^, full-quantile normalization^46^, and relative log-expression scaling normalization^47^.

*Linnorm*^29^ performs a prior logarithmic transformation on the expression data, and the dataset is fitted to a linear model that does not need to go through the origin. This allows expression level to be adjusted both linearly and exponentially. Bioconductor R package *linnorm*^29^ were used with the default settings.

*Scran*^30^ computes the scaling factors on pooled expression measures and then deconvolved to obtain cell-specific factors. The method is implemented in the Bioconductor R package *scran*^30^. Pool sizes from 20 to the 100 (intervals of five) were considered.

*Seurat*^27^ divides the transcript counts for each cell by the total counts for that cell and multiplied by the scale factor (i.e., default scaling factor is 10000) followed by natural-log transformation. *Seurat* is implement in the Bioconductor R package *Seurat*^27^.

*Signac*^41^ is an extension of Seurat for the normalization and analysis of single-cell chromatin datasets. It computes term frequency-inverse document frequency (TF-IDF) normalization of the peak matrix by dividing the accessibility of each peak in a cell by the cell’s total accessibility and multiplying this by the inverse accessibility of the peak in the cell population. This TF-IDF matrix is then log-transformed^27^.

## Data and code availability

The scPSD method has been implemented in MATLAB (https://github.com/VafaeeLab/psdMAT), Python (https://github.com/VafaeeLab/psdPy) and R package (https://github.com/VafaeeLab/psdR). To assure the reproducibility of the reported results, the data and pipeline developed for this study are available at (https://github.com/VafaeeLab/psdMAT).

## Acknowledgement

This research was funded by the UNSW Cellular Genomics Futures Institute, University of New South Wales, Sydney Australia. We also acknowledge UNSW Research Technology (ResTech) for providing high-performance computing resources (Katana) enabling extensive analysis conducted in this study. We also acknowledge constructive comments from Dr Omid Faridani.

## Author contributions

FV conceived and led the study and guided the method development. SMZ developed the scPSD method and the corresponding MATLAB package. SMZ and FV conducted the analyses and produced the results. FV and SMZ wrote the manuscript and generated display items. DGO and FVM curated the public datasets and provided input to method evaluation. FK performed DR analyses and developed the Python implementation of scPSD. AV conducted the TI analyses and developed the scPSD R package. FZ curated in-house scRNA-seq datasets and provided input on method evaluation. All authors critically reviewed the manuscript and approved its final version.

## Supplementary Tables

**Table 1.** Summary of scRNA-seq datasets used for the evaluation of scPSD.

| # | Dataset | Protocol | Accession | Species | Tissue type | Cells | Genes | Cell Types | Min cell type % | Max cell type % | UMI/Read | Cell type determination* |
| --- | --- | --- | --- | --- | --- | --- | --- | --- | --- | --- | --- | --- |
| 1 | chuBatch1<sup>1</sup> | SMARTer | GSE75748 | human | stem cells | 350 | 19097 | 5 | 14 | 24.86 | Read | Gold |
| 2 | chuBatch2<sup>1</sup> | SMARTer | GSE75748 | human | stem cells | 425 | 19097 | 6 | 9.65 | 20.47 | Read | Gold |
| 3 | Linnarsson<sup>2</sup> (cerebellum) | Chromium | SRP135960 | mouse | brain | 12312 | 17984 | 8 | 0.48 | 64.75 | UMI | Prediction |
| 4 | Colon FACS<sup>3</sup> | Smart-Seq2 | GSE132042 | mouse | colon | 4149 | 23433 | 6 | 0.6 | 19.62 | Read | Silver |
| 5 | Fat FACS<sup>3</sup> | Smart-Seq2 | GSE132042 | mouse | fat | 5862 | 23433 | 11 | 0.46 | 3.77 | Read | Silver |
| 6 | Heart 10X<sup>3</sup> | Chromium | GSE132042 | mouse | heart | 654 | 23433 | 7 | 3.21 | 33.94 | UMI | Prediction |
| 7 | Heart FACS<sup>3</sup> | Smart-Seq2 | GSE132042 | mouse | heart | 7115 | 23433 | 11 | 0.16 | 17.95 | Read | Prediction |
| 8 | Kidney FACS<sup>3</sup> | Smart-Seq2 | GSE132042 | mouse | kidney | 865 | 23433 | 7 | 4.39 | 7.51 | Read | Prediction |
| 9 | Liver FACS<sup>3</sup> | Smart-Seq2 | GSE132042 | mouse | liver | 981 | 23433 | 6 | 2.96 | 19.98 | Read | Prediction |
| 10 | Lung FACS<sup>3</sup> | Smart-Seq2 | GSE132042 | mouse | lung | 1923 | 23433 | 17 | 0.1 | 0.68 | Read | Silver |
| 11 | Pancreas FACS<sup>3</sup> | Smart-Seq2 | GSE132042 | mouse | pancreas | 1961 | 23433 | 10 | 1.48 | 20.96 | Read | Prediction |
| 12 | baron mouse<sup>4</sup> | inDrop | GSE84133 | mouse | pancreas | 1886 | 14878 | 13 | 0.32 | 11.56 | UMI | Prediction |
| 13 | deng reads<sup>5</sup> | Smart-Seq2 | GSE45719 | mouse | embryo | 268 | 22431 | 6 | 4.48 | 13.81 | Read | Gold |
| 14 | fan<sup>6</sup> | SUPeR-seq | GSE53386 | mouse | embryo | 66 | 26357 | 6 | 10.61 | 33.33 | Read | Gold |
| 15 | goolam<sup>7</sup> | Smart-Seq2 | E-MTAB-3321 | mouse | embryo | 124 | 41480 | 5 | 4.84 | 25.81 | Read | Gold |
| 16 | klein<sup>8</sup> | inDrop | GSE65525 | mouse | embryo stem cells | 2717 | 24175 | 4 | 11.15 | 34.34 | UMI | Gold |
| 17 | li<sup>9</sup> | SMARTer | GSE81861 | human | colorectal tumor | 561 | 55186 | 9 | 4.1 | 12.3 | Read | Prediction |
| 18 | manno human<sup>10</sup> | STRT-Seq UMI | GSE76381 | human | brain | 4029 | 20560 | 56 | 0.12 | 0.45 | UMI | Prediction |
| 19 | manno mouse<sup>10</sup> | STRT-Seq UMI | GSE76381 | mouse | brain | 2150 | 24378 | 32 | 0.56 | 1.26 | UMI | Prediction |
| 20 | marques<sup>11</sup> | C1 | GSE75330 | mouse | brain | 5053 | 23556 | 13 | 1.5 | 9.12 | Read | Prediction |
| 21 | pollen<sup>12</sup> | SMARTer | SRP041736 | human | neonatal-foreskin | 301 | 23730 | 11 | 2.33 | 17.94 | Read | Prediction |
| 22 | tasic reads<sup>13</sup> | SMARTer | GSE71585 | mouse | brain | 1679 | 24150 | 18 | 0.54 | 16.38 | Read | Silver |
| 23 | yan<sup>14</sup> | Tang | GSE36552 | human | embryo | 90 | 20214 | 6 | 6.67 | 17.78 | Read | Gold |
| 24 | zhengmix4eq<sup>15</sup> | GemCode | SRP073767 | human | immune | 3994 | 15568 | 4 | 3.05 | 8.55 | UMI | Gold |
| 25 | zhengmix4uneq<sup>15</sup> | GemCode | SRP073767 | human | immune | 6498 | 16443 | 4 | 7.69 | 46.14 | UMI | Gold |

**Table 2.** Summary of scRNA-seq in-house datasets used for the evaluation of scPSD.

| Dataset | Accession | Tissue type | Cells | Genes | # of Cell Types | Cell Types (#) |
| --- | --- | --- | --- | --- | --- | --- |
| Domingo-Gonzalez and Zanini et al.<sup>16</sup> | GSE147668 | mouse murine lung **immune** cells | 5234 | 18072 | 16 | Mac I (737), Mac II (391), Mac III (563), Mac IV (332), Mac V (847), mastCell (67), mastCell II (22), basophil (102), TCell (418), NKCell (111), BCell (755), ILCell (60), DC I (94), DC II (67), DC III (36), neutrophil (632). |
| Zanini et al.<sup>17</sup> | GSE172251 | mouse murine lung **mesenchymal** cells | 5479 | 18072 | 16 | Fibroblast Precursor (764), Proliferating Myofibroblast (118), Early Alveolar Fibroblast (1618), Alveolar Fibroblast (250), Early Adventitial Fibroblast (598), Adventitial Fibroblast (241), Male Hyperoxic Fibroblast (55), Myofibroblast And Smooth Muscle Precursor (241), Early Airway Smooth Muscle (285), Airway Smooth Muscle (113), Myofibroblast (520), Proliferating Fibroblast (157), Vascular Smooth Muscle (201), Pericyte (281), Proliferating Pericyte (19), Striated Muscle (18) |
| Zanini et al.<sup>18</sup> | GSE159804 | mouse murine lung **endothelial** cells | 2930 | 18072 | 10 | Arterial EC I (96), Arterial EC II (42), Venous EC (203), Lymphatic EC (82), Proliferative Venous EC (10), Proliferative EC (390), Early Car4 Capillaries (840), Late Car 4 Capillaries (622), Car 4 Capillaries (332), Nonproliferative Embryonic EC (313) |

**Table 3.** Cell-specific SVM classification performance (sensitivity and specificity over test set, i.e., 20% random holdout cells) using datasets detailed in Supplementary Table 2, before and after scPSD feature extraction (CPM normalization).

|  | Cell type | Before scPSD |  | After scPSD |  |
| --- | --- | --- | --- | --- | --- |
|  |  | Sensitivity | Specificity | Sensitivity | Specificity |
| Immune | Mac I | 0.979 | 0.889 | 0.950 | 0.986 |
|  | Mac II | 0.0 | 0.967 | 0.848 | 0.988 |
|  | Mac III | 0.915 | 0.958 | 1.0 | 0.989 |
|  | Mac IV | 0.418 | 0.960 | 0.896 | 0.995 |
|  | Mac V | 0.848 | 0.8644 | 0.966 | 0.990 |
|  | mastCell | 0.882 | 0.964 | 1.0 | 0.995 |
|  | mastCell II | 0.0 | 0.966 | 0.75 | 0.995 |
|  | basophil | 0.0 | 0.963 | 1.0 | 0.995 |
|  | TCell | 0.825 | 0.942 | 0.950 | 0.990 |
|  | NKCell | 0.0 | 0.963 | 0.905 | 0.994 |
|  | BCell | 0.942 | 0.955 | 0.985 | 0.992 |
|  | ILCell | 0.0 | 0.966 | 0.929 | 0.992 |
|  | DC I | 0.0 | 0.964 | 1.0 | 0.995 |
|  | DC II | 0.0 | 0.963 | 0.917 | 0.991 |
|  | DC III | 0.0 | 0.963 | 1.0 | 0.995 |
|  | neutrophil | 0.898 | 0.887 | 0.977 | 0.997 |
| Mesenchymal | Fibroblast Precursor | 0.604 | 0.803 | 0.968 | 0.991 |
|  | Proliferating Myofibroblast | 0.217 | 0.846 | 0.880 | 0.995 |
|  | Early Alveolar Fibroblast | 0.861 | 0.622 | 0.967 | 0.979 |
|  | Alveolar Fibroblast | 0.477 | 0.847 | 0.955 | 0.995 |
|  | Early Adventitial Fibroblast | 0.636 | 0.803 | 0.926 | 0.988 |
|  | Adventitial Fibroblast | 0.0 | 0.856 | 0.887 | 0.997 |
|  | Male Hyperoxic Fibroblast | 0.0 | 0.847 | 0.846 | 0.996 |
|  | Myofibroblast & Smooth Muscle Precursor | 0.607 | 0.844 | 0.911 | 0.994 |
|  | Early Airway Smooth Muscle | 0.603 | 0.842 | 0.956 | 0.989 |
|  | Airway Smooth Muscle | 0.0 | 0.852 | 1.0 | 0.997 |
|  | Myofibroblast | 0.468 | 0.855 | 0.863 | 0.991 |
|  | Proliferating Fibroblast | 0.217 | 0.846 | 0.870 | 0.995 |
|  | Vascular Smooth Muscle | 0.026 | 0.851 | 0.921 | 0.996 |
|  | Pericyte | 0.754 | 0.829 | 0.983 | 0.995 |
|  | Proliferating Pericyte | 0.0 | 0.848 | 1.0 | 0.997 |
|  | Striated Muscle | 0.0 | 0.851 | 1.0 | 0.997 |
| Endothelial | Arterial EC I | 0.0 | 0.942 | 0.5 | 0.989 |
|  | Arterial EC II | 0.0 | 0.945 | 1.0 | 0.995 |
|  | Venous EC | 0.0 | 0.942 | 0.844 | 0.990 |
|  | Lymphatic EC | 0.0 | 0.948 | 0.947 | 0.998 |
|  | Proliferative Venous EC | 0.0 | 0.944 | 0.75 | 0.998 |
|  | Proliferative EC | 0.870 | 0.914 | 0.961 | 0.987 |
|  | Early Car4 Capillaries | 0.946 | 0.420 | 0.912 | 0.956 |
|  | Late Car 4 Capillaries | 0.307 | 0.907 | 0.921 | 0.989 |
|  | Car 4 Capillaries | 0.552 | 0.947 | 0.940 | 0.992 |
|  | Nonproliferative Embryonic EC | 0.0 | 0.940 | 0.9 | 0.979 |

**Table 4.** Summary of dimensionality reduction methods studied in Supplementary Figure X.

| Method | Abbrev. | Brief Description | Package |
| --- | --- | --- | --- |
| Bayesian decomposition<sup>19</sup> | BD | Uses a Gibbs sampler to estimate the posterior density of NMF factors | nimfa |
| Factor Analysis<sup>20</sup> | FA | Linear generative model with Gaussian latent variables | sklearn |
| Fast Independent Component Analysis<sup>21</sup> | FICA | Extracts statistically independent non-Gaussian signals | sklearn |
| Gaussian Random Projection<sup>22</sup> | GRP | Projecting the original input space on a randomly generated matrix where components are drawn from a normal distribution | sklearn |
| Incremental PCA<sup>23</sup> | IPCA | A mini-batch implementation of PCA | sklearn |
| Isometric Mapping<sup>24</sup> | ISOMAP | An extension on MDS that attempts to preserve geodesic distances | sklearn |
| Iterated Conditional Modes<sup>19</sup> | ICM | Like BD, but uses modes in place of random samples in the Gibbs sampler | nimfa |
| ivis<sup>25</sup> | IVIS | Uses Siamese neural networks to embed data | ivis |
| Kernel PCA<sup>26</sup> (with cosine, polynomial, and radial basis function transformations) | KPCA-(COS/POL/RBF) | Utilizes the kernel trick to allow for non-linear representations of the data. | sklearn |
| Latent Dirichlet allocation<sup>27</sup> | LDA | Bayesian generative model utilizing a Dirichlet prior over latent space | sklearn |
| Least squares NMF<sup>28</sup> | LSNMF | Alternative to multiplicative NMF which claims to have faster convergence | nimfa |
| Locally linear embedding<sup>29</sup> | LLE | Similar to isomap, but attempts to preserve local distances | sklearn |
| Non-negative Matrix Factorization<sup>30,31</sup> | NMF-(NNSVD/LEE) | Factorizes a (non-negative) data into two non-negative matrices. | sklearn/nimfa |
| Non-smooth NMF (nsnmf)<sup>32</sup> | NSNMF | Applies smoothing kernel into the factorization step reducing sparsity | nimfa |
| Potential of Heat-diffusion for Affinity-based Trajectory Embedding<sup>33</sup> | PHATE | Utilizes a heat diffusion model to preserve branching structures in data | phate |
| Principle Component Analysis<sup>34</sup> | PCA | Construct a low dimensional, orthogonal basis that maximally explains observed variance in the data. | sklearn |
| Probabilistic non-negative Matrix Factorization<sup>35</sup> | PMF | Assumption that data follows a multinomial distribution and performs expectation maximization | nimfa |
| Probabilistic sparse MF<sup>36</sup> | PSMF | Assumption of Gaussian noise within the data, and sparsity constraint added for factorization | nimfa |
| Sparse Autoencoder for Clustering, Imputing, and Embedding<sup>37</sup> | SAUCIE | Sparse Autoencoder Neural Network | saucie |
| Sparse nmf<sup>38</sup> | SNMF | Addition of sparsity constraint to factorization process | nimfa |
| sparse PCA<sup>39</sup> | SPCA(-BATCH) | An extension of PCA which aims to produce sparse (with many zeros) components | sklearn |
| Sparse Random Projection<sup>40</sup> | SRP | Similar to GRP, but matrix elements are drawn from a zero-inflated, centered uniform distribution. | sklearn |
| Laplacian eigenmaps<sup>41</sup> | SPECTRAL | Attempts to reconstruct the manifold in low dimensional space by preserving the distances of the nearest neighbors to each point | sklearn |
| t-distributed Stochastic Neighbour Embedding<sup>42</sup> | TSNE | Minimizes the KL divergence of the estimated distribution of points in high dimensional points with a similar low-dimensional distribution. | Mutlicore TSNE |
| Truncated Singular Value Decomposition<sup>34</sup> | TSVD | PCA without centring and scaling | sklearn |
| Uniform Manifold Approximation and Projection<sup>43</sup> | UMAP | Constructs a fuzzy topological representation of the data to preserve in the low dimensional embedding | Umap-learn |
| Variational autoencoder<sup>44</sup> | VASC | Uses neural network to map latent variables to zero-inflated negative binomial distribution | vasc |
| Variational projection<sup>45</sup> | VPAC | Uses a variational projection algorithm to infer a Gaussian mixture in latent space | vpac |
| Zero-inflated Factor Analysis<sup>46</sup> | ZIFA | Uses a latent variable model, similar to FA, in which is modulated by the addition of a zero-inflated layer meant to simulate dropout. | ZIFA |

**Table 5.**
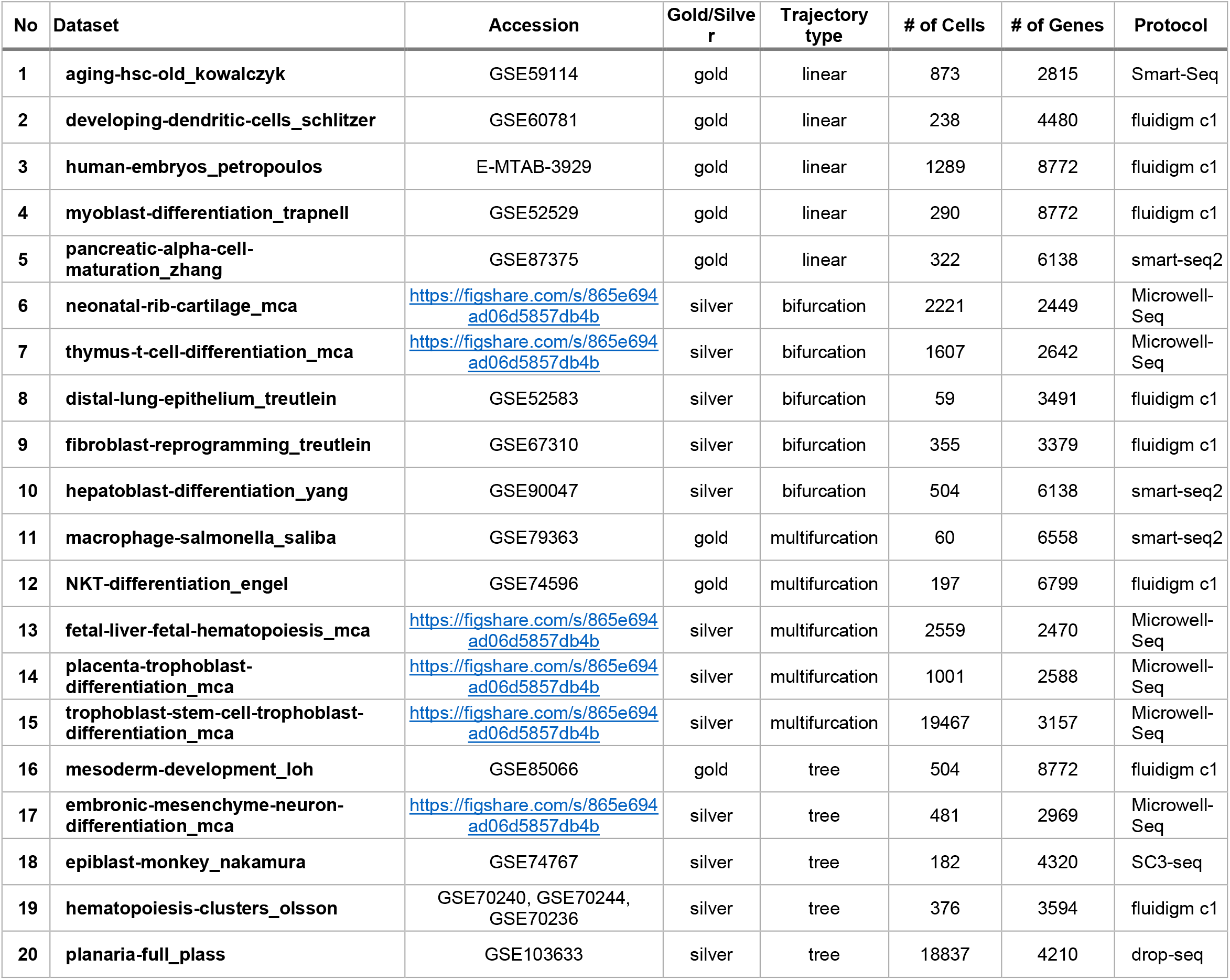
*Summary of trajectory inference (TI) datasets used for the evaluation of scPSD*. The R package suite created alongside the TI benchmarking study by Saelens et al47 (https://github.com/dynverse) was used for performing the trajectory inference. Datasets uploaded to zenodo (https://zenodo.org/record/1443566) as part of the same study was used here; datasets were classified by Saelens *et al* as ‘gold standard’ if the reference trajectory was not extracted from the expression data itself, such as via cellular sorting or cell mixing or as ‘silver standard’, otherwise. Please refer to the original paper for further details on datasets.

**Table 6.**
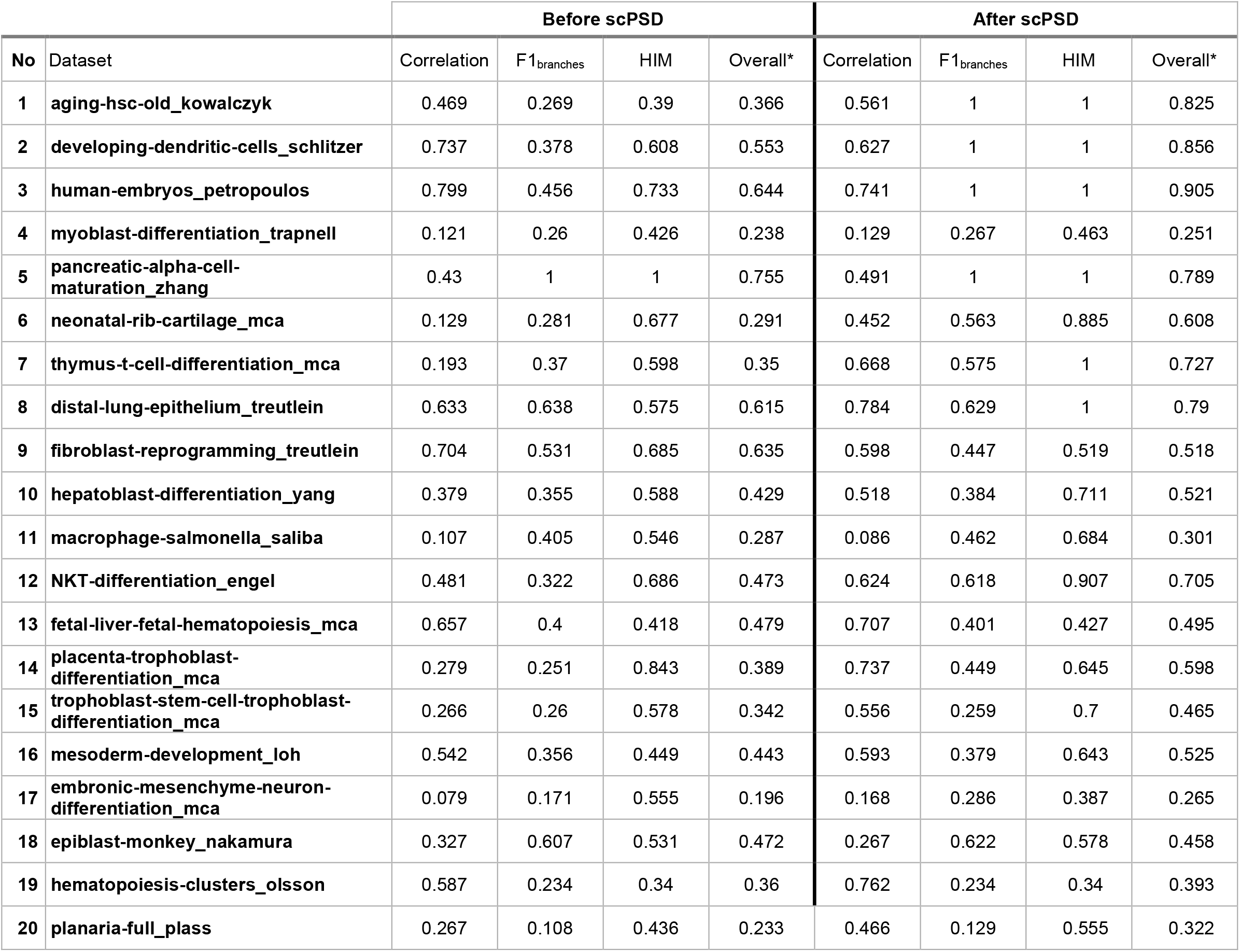
Details of metrics comparing the performance of MST trajectory inference before and after scPSD transformation. * Last column represents the overall score computed as the geometric mean of the other three metrics.

---

**Supplementary Fig 1.**
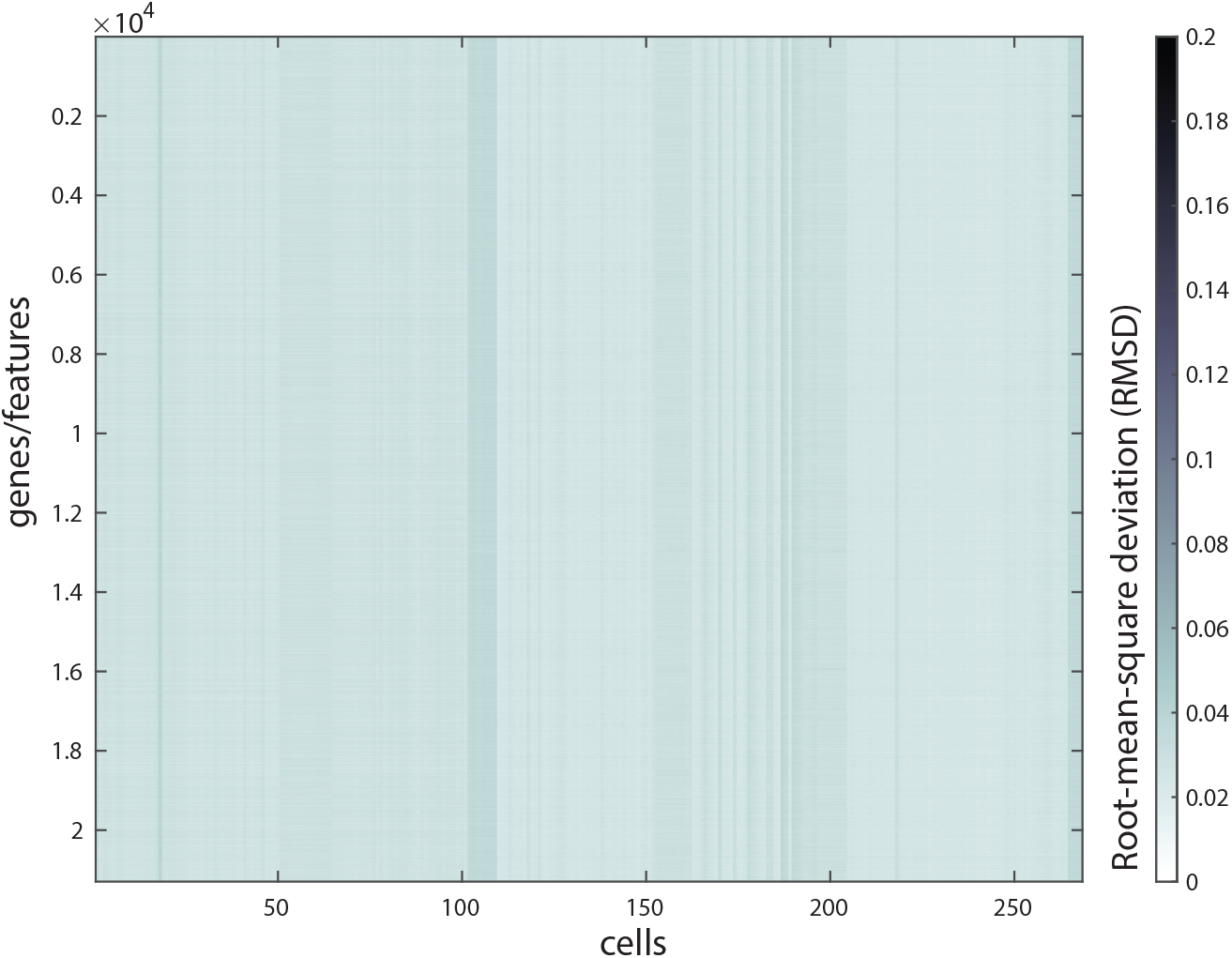
The Heatmap shows root-mean-square deviations (RMSD) of scPSD-transformed gene expression after 100 random shuffles using deng reads dataset (database # 13 in Supplementary Table 1 with n = 22,431 genes/features and m = 268 cells). In order to show independency of transformation on initial ordering of features, a gene ordering was randomly picked as the ‘reference order’ and then the order of transcripts was shuffled 100 times prior to scPSD transformation. After applying scPSD, the transformed matrices were rearranged to unify gene ordering based on the ‘reference order’. Accordingly, for each gene k in cell j, RMSD was calculated estimating the mean-square deviation of transformed gene expression compared to the corresponding expression in the reference matrix. The heatmap is therefore an nxm matrix where each element shows RMSD for individual genes k = 1, …, n across cells j = 1, … m. The minimum and maximum RMSD obtained are 2.47e-18 and 4.33e-2, respectively.

**Supplementary Fig 2.**
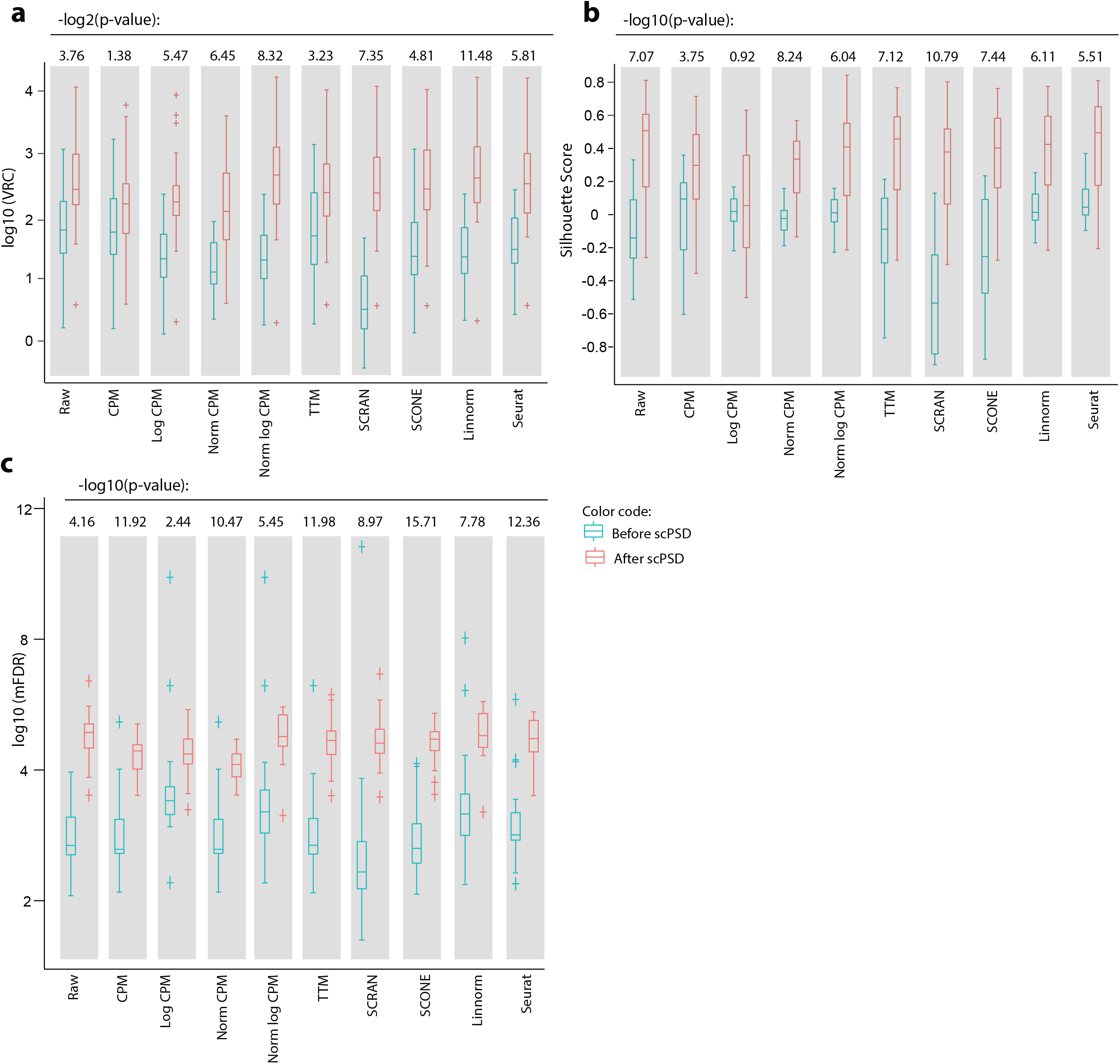
Boxplots representing the distribution of Variance Ratio Criterion, VRC (a), Silhouette Score, SS (b), and multiclass Fisher’s Discriminant Ratio, mFDR (C) across 25 scRNA-seq datasets (c.f. Supplementary Table 1) before (green) and after (red) scPSD transformations. The statistical significance of improvements on metrics upon ^2^ scPSD transformation has been assessed using t-test; -log10(p-values) are reported on top of each pairwise comparison.

**Supplementary Fig 3.**
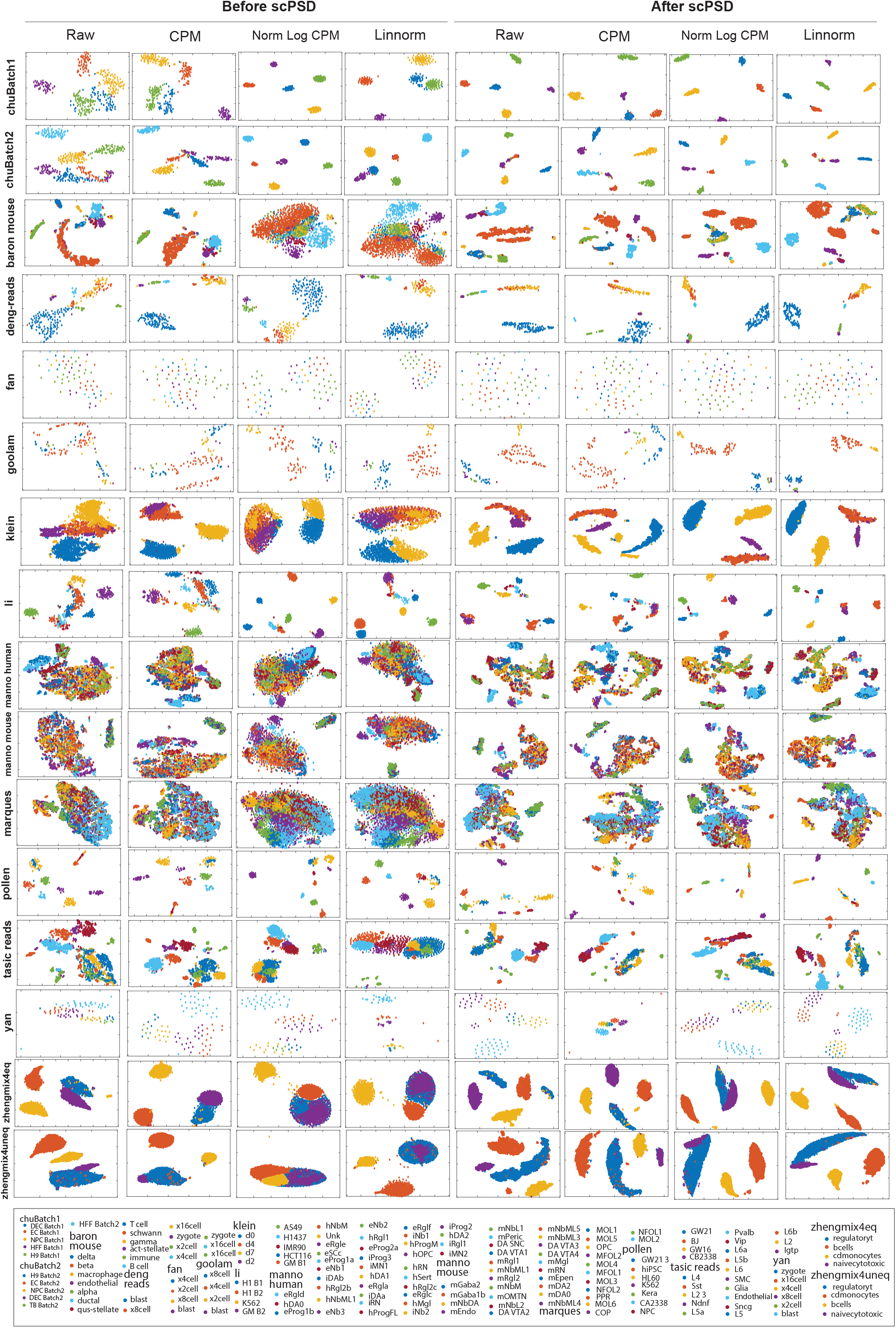
2D visualization (t-SNE dimensionality reduction) of scRNA-seq datasets (c.f. Supplementary Table 1) demonstrating visual separation of cell types upon scPSD transformation of raw or normalised data.

**Supplementary Fig 4.**
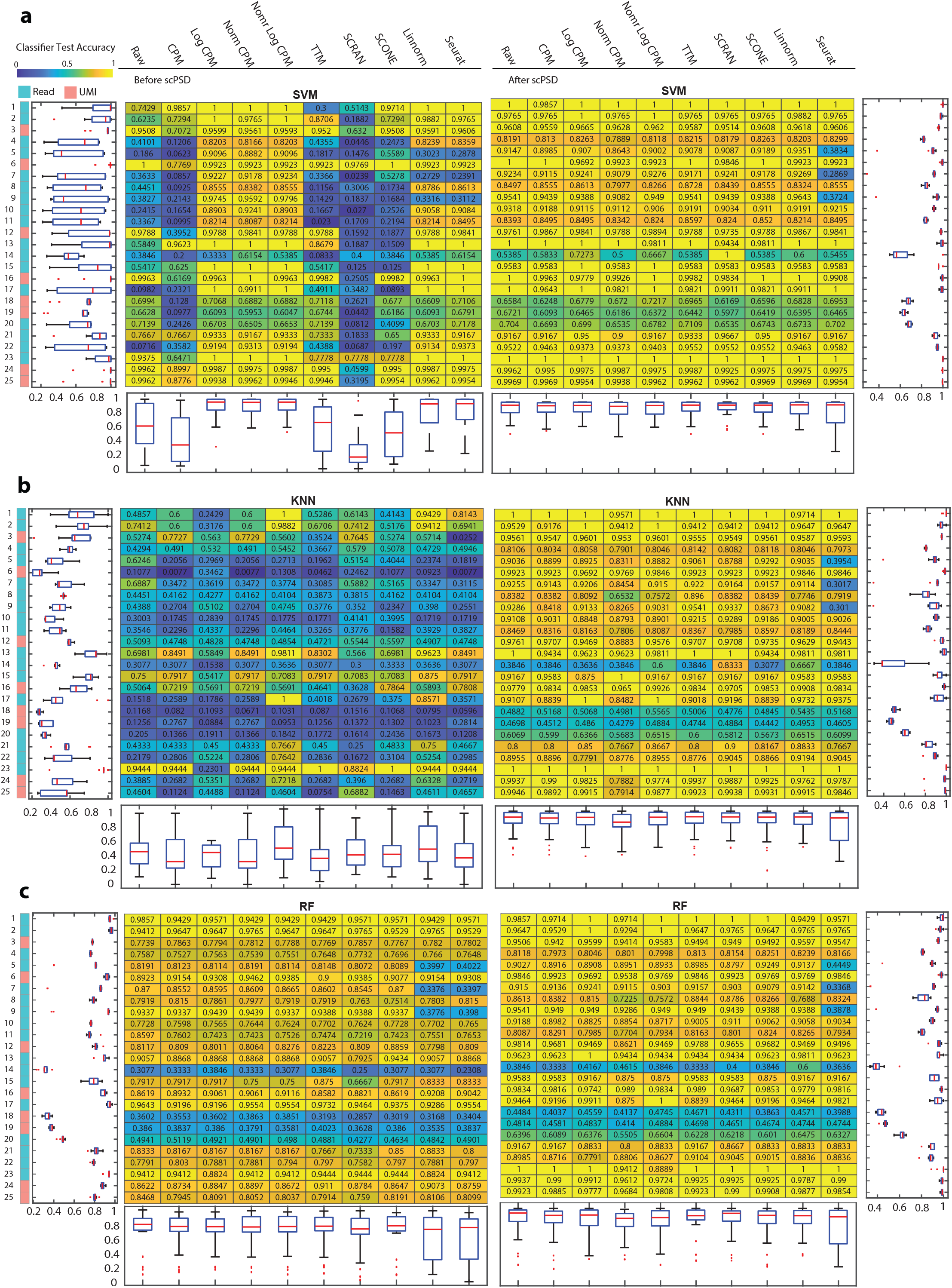
Heatmaps with marginal boxplots representing cell type classification performance across each dataset before and after scPSD feature extraction (with different normalization methods used as upstream processing). Three classifiers were studied including support vector machine, SVM (a), random forest, RF (b) and k-nearest neighbor, KNN (C). For each dataset, the performance was evaluated based on the classification accuracy over a holdout test set (20% random split of a dataset)

**Supplementary Fig 4.**
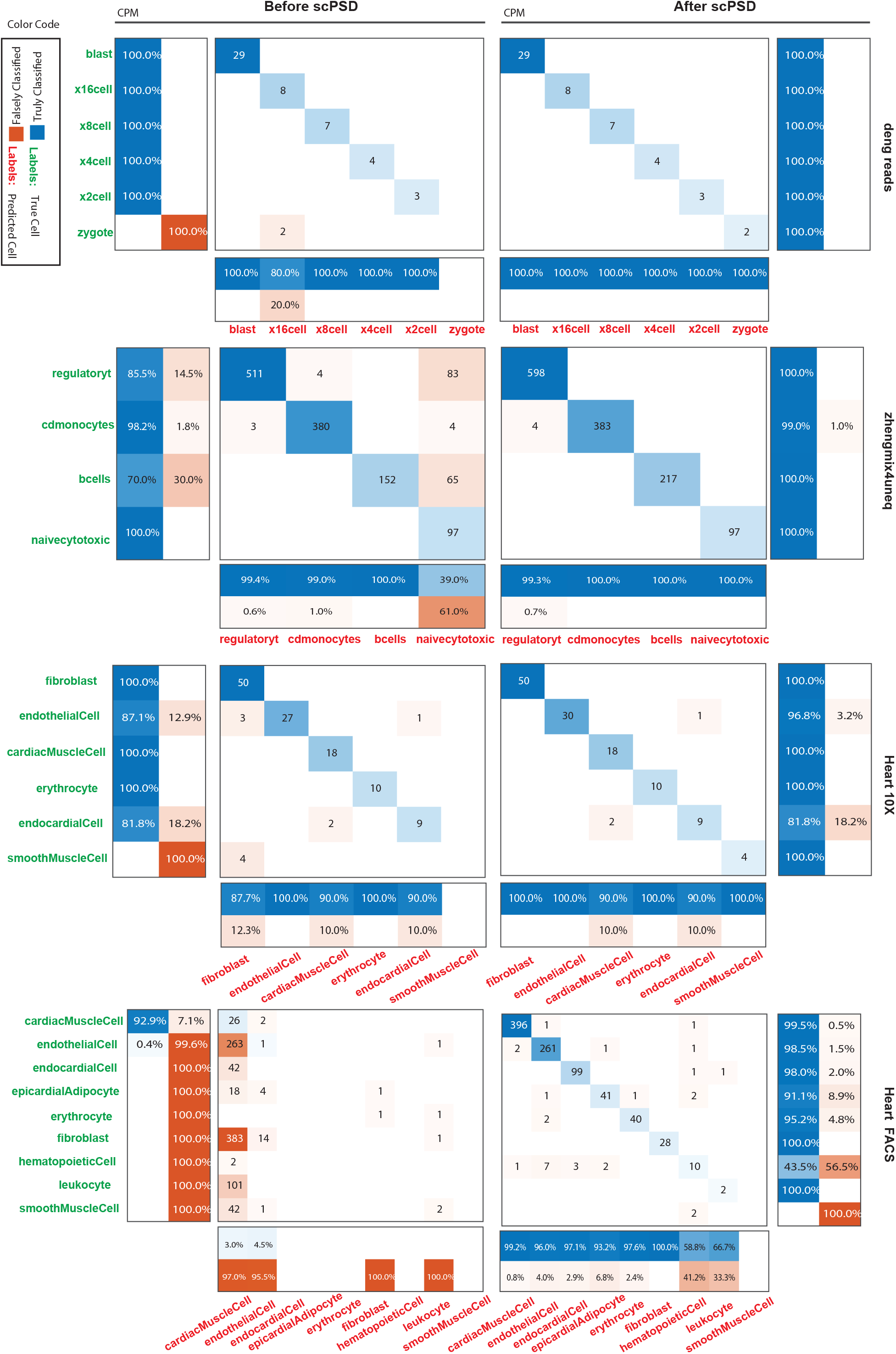
Confusion matrices (CMs) detailing the performance of SVM classification on test data (randomly picked 20% of dataset). Each row of a matrix represents the instances in an actual class (green, determined based on the dataset annotations) while each column represents the instances in a predicted class (red, determined by trained SVM classifier). CMs were visualised for four datasets with comparatively reliable annotations (c.f. Supplementary Table 1) before and after scPSD transformation to assess classification performance on cell types with small populations (cell types or classes are sorted in descending order of their size). The results demonstrate the capacity of scPSD to enhance rare cell type identification. Similar results were observed on other public and in-house datasets (c.f. Fig 2).

**Supplementary Fig 8.**
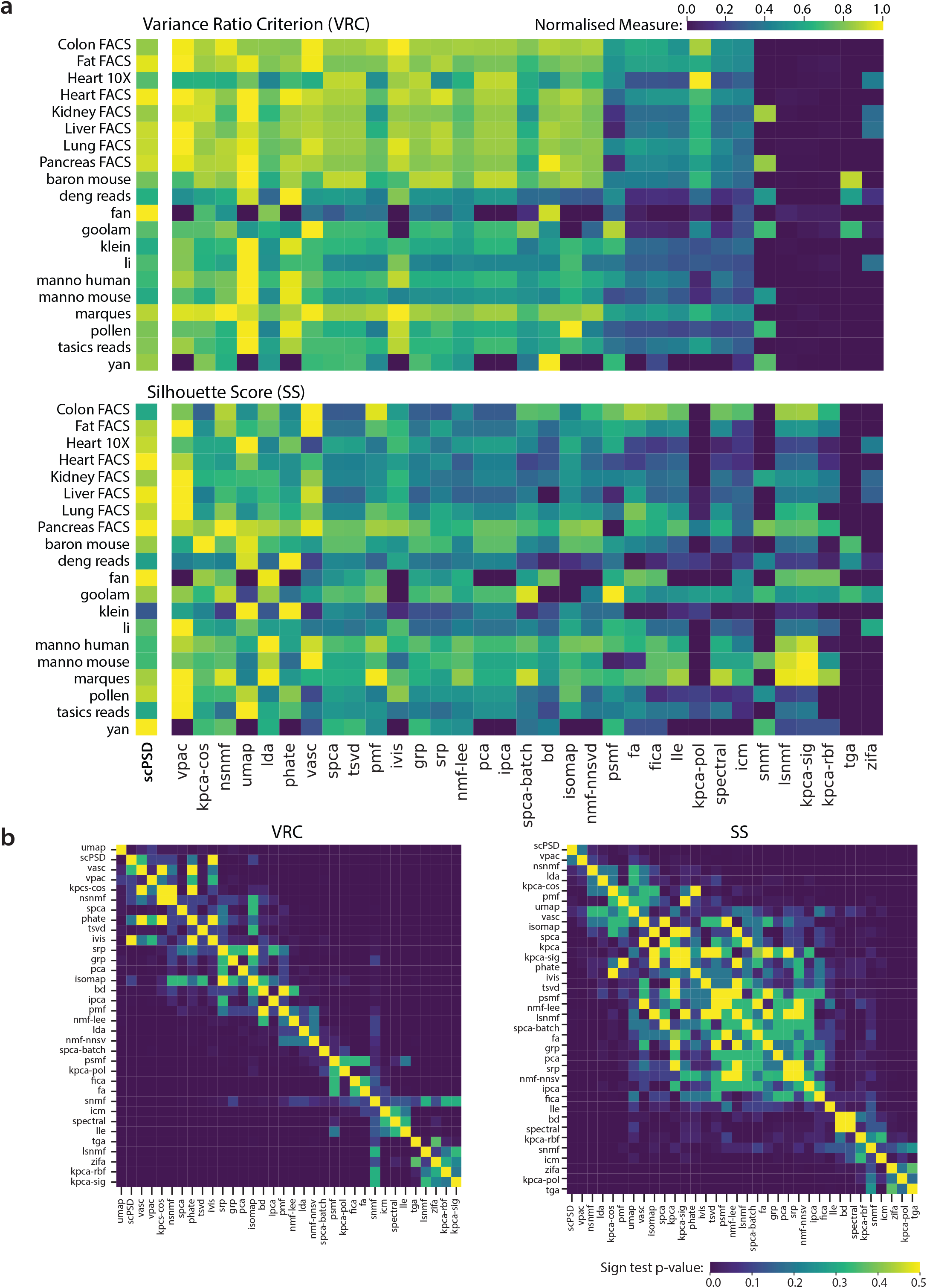
**(a)** Heatmaps of the SS with Euclidean distance and VRC for 33 dimensionality reduction methods with embedding size of 96 (methods detailed in Supplementary Table 2). Values were standardized using min-max normalization applied per-dataset (Supplementary Table 1). Methods are orderd from left to right based on their average VRC score across all databases; scPSD shows the highest average VRC and placed at the left-most position. **(b)** Heatmaps of p-values from pairwise sign-tests comparing measures for each method across all datasets. It depicts the relative ranking of dimensionality reduction methods, as well as providing information as to which methods were roughly equivalent.

**Supplementary Fig 7.**
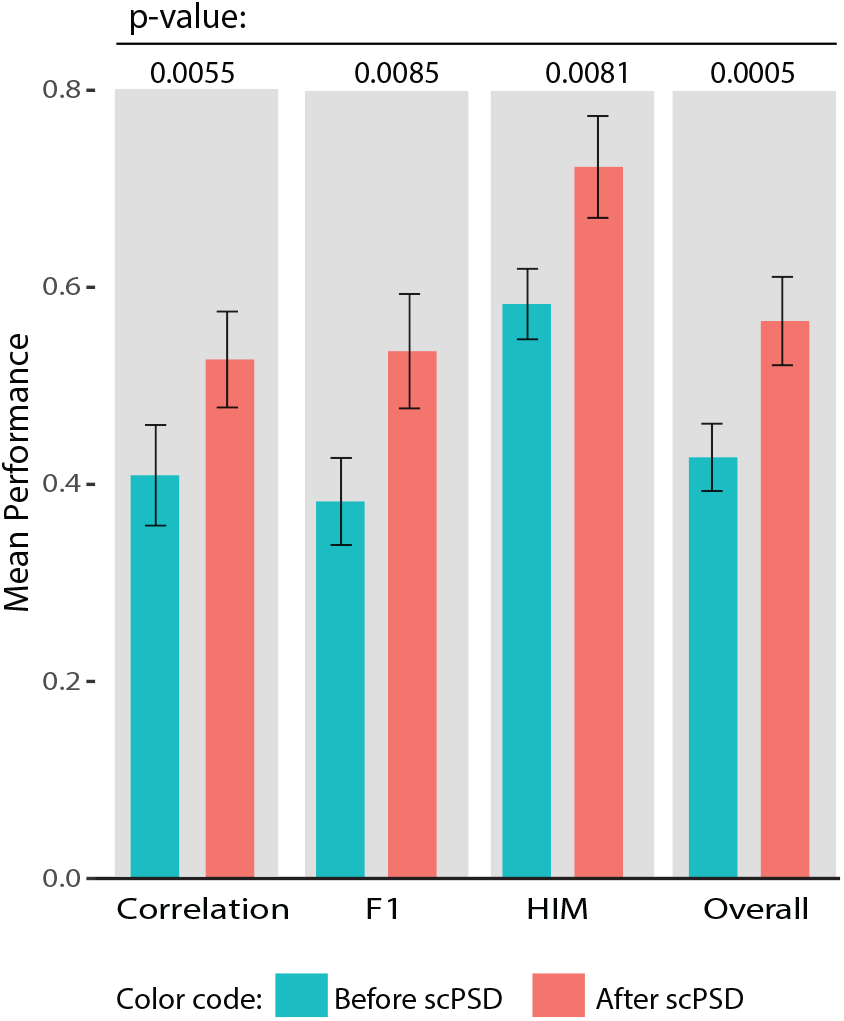
Trajectory inference (Tl) analysis. Bar plots representing the mean performance of minimum spanning tree (MST) in infer-ring the topology of 20 datasets (Supplementary Table 5) representing trajectories with different topologies (linear, bifurcation, multifurcation, and tree). The error bars represent the standard error. MST method, studied by (Saelens et al, Nature Biotechnology, 2019), is a high-performing Off-the-shelf method which performs PCA dimensionality reduction, followed by clustering using the R mclust package, and finally connects clusters using a minimum spanning tree algorithm. The performance metrics include 1) HIM which measures the topological similarity 2) F1 between branch assignments, 3) correlation between geodesic distances, and 4) overall score computed as geometric mean of the three metrics (c.f. Saelens et al for details of metrics). P-value of paired t-test on top of the plot shows the significance of comparison before vs after scPSD transformation. for each metric. This initial result is not generalisable to any TI method and the effect of scPSD transformation on other well-preforming TI methods requires further investigation.

## References

1. Zhu, C., Preissl, S. & Ren, B.J.N.m. Single-cell multimodal omics: the power of many. Nature methods 17, 11–14 (2020).

2. Lähnemann, D. et al. Eleven grand challenges in single-cell data science. Genome biology 21, 1–35 (2020).

3. Chen, H. et al. Assessment of computational methods for the analysis of single-cell ATAC-seq data. Genome biology 20, 1–25 (2019).

4. Patruno, L. et al. A review of computational strategies for denoising and imputation of single-cell transcriptomic data. Briefings in Bioinformatics (2020).

5. Raimundo, F., Papaxanthos, L., Vallot, C. & Vert, J.-P.J.C.O.i.S.B. Machine learning for single cell genomics data analysis. Current Opinion in Systems Biology (2021).

6. Bonidia, R.P. et al. Feature extraction approaches for biological sequences: a comparative study of mathematical features. Briefings in Bioinformatics (2020).

7. Tsuyuzaki, K., Sato, H., Sato, K. & Nikaido, I.J.G.b. Benchmarking principal component analysis for large-scale single-cell RNA-sequencing. Genome biology 21, 1–17 (2020).

8. Van der Maaten, L. & Hinton, G.J.J.o.m.l.r. Visualizing data using t-SNE. Journal of machine learning research 9(2008).

9. Becht, E. et al. Dimensionality reduction for visualizing single-cell data using UMAP. Nature biotechnology 37, 38–44 (2019).

10. Pierson, E. & Yau, C.J.G.b. ZIFA: Dimensionality reduction for zero-inflated single-cell gene expression analysis. Genome biology 16, 1–10 (2015).

11. Risso, D., Perraudeau, F., Gribkova, S., Dudoit, S. & Vert, J.-P.J.N.c. A general and flexible method for signal extraction from single-cell RNA-seq data. Nature communications 9, 1–17 (2018).

12. Lopez, R., Regier, J., Cole, M.B., Jordan, M.I. & Yosef, N.J.N.m. Deep generative modeling for single-cell transcriptomics. Nature methods 15, 1053–1058 (2018).

13. Xiong, L. et al. SCALE method for single-cell ATAC-seq analysis via latent feature extraction. Nature communications 10, 1–10 (2019).

14. Koch, F.C., Sutton, G.J., Voineagu, I. & Vafaee, F.J.b. Supervised Application of Internal Validation Measures to Benchmark Dimensionality Reduction Methods in scRNA-seq Data. Briefings in Bioinformatics bbab304(2020).

15. Sun, S., Zhu, J., Ma, Y. & Zhou, X.J.G.b. Accuracy, robustness and scalability of dimensionality reduction methods for single-cell RNA-seq analysis. Genome biology 20, 1–21 (2019).

16. Stoica, P. & Moses, R.L. Spectral analysis of signals. (2005).

17. Cochran, W.T. et al. What is the fast Fourier transform? J Proceedings of the IEEE 55, 1664–1674 (1967).

18. Cover, T.M. Elements of information theory, (John Wiley & Sons, 1999).

19. Chanda, P. et al. Information Theory in Computational Biology: Where We Stand Today. Entropy 22, 627 (2020).

20. Vinga, S.J.B.i.b. Information theory applications for biological sequence analysis. Briefings in bioinformatics 15, 376–389 (2014).

21. Koch, F.C., Sutton, G.J., Voineagu, I. & Vafaee, F.J.b. Supervised Application of Internal Validation Measures to Benchmark Dimensionality Reduction Methods in scRNA-seq Data. bioRxiv (2020).

22. Rousseeuw, P.J.J.J.o.c. & mathematics, a. Silhouettes: a graphical aid to the interpretation and validation of cluster analysis. J Journal of computational applied mathematics 20, 53–65 (1987).

23. Calinski, T., Harabasz, J.J.C.i.S.-t. & Methods. A dendrite method for cluster analysis. J Communications in Statistics-theory Methods 3, 1–27 (1974).

24. Webb, A.R. Statistical pattern recognition, (John Wiley & Sons, 2003).

25. Robinson, M.D. & Oshlack, A. A scaling normalization method for differential expression analysis of RNA-seq data. Genome Biol 11, R25 (2010).

26. Robinson, M.D., McCarthy, D.J. & Smyth, G.K. edgeR: a Bioconductor package for differential expression analysis of digital gene expression data. Bioinformatics 26, 139–40 (2010).

27. Stuart, T. et al. Comprehensive integration of single-cell data. Cell 177, 1888-1902. e21 (2019).

28. Cole, M.B. et al. Performance Assessment and Selection of Normalization Procedures for Single-Cell RNA-Seq. Cell Syst 8, 315–328 e8 (2019).

29. Yip, S.H., Wang, P., Kocher, J.-P.A., Sham, P.C. & Wang, J. Linnorm: improved statistical analysis for single cell RNA-seq expression data. Nucleic acids research 45, e179–e179 (2017).

30. Lun, A.T., Bach, K. & Marioni, J.C. Pooling across cells to normalize single-cell RNA sequencing data with many zero counts. Genome Biol 17, 75 (2016).

31. Abdelaal, T. et al. A comparison of automatic cell identification methods for single-cell RNA sequencing data. Genome biology 20, 1–19 (2019).

32. Jiang, L., Chen, H., Pinello, L. & Yuan, G.-C.J.G.b. GiniClust: detecting rare cell types from single-cell gene expression data with Gini index. Genome biology 17, 1–13 (2016).

33. Deng, Q., Ramsköld, D., Reinius, B. & Sandberg, R.J.S. Single-cell RNA-seq reveals dynamic, random monoallelic gene expression in mammalian cells. Science 343, 193–196 (2014).

34. Saelens, W., Cannoodt, R., Todorov, H. & Saeys, Y.J. A comparison of single-cell trajectory inference methods. Nature biotechnology 37, 547–554 (2019).

35. Jurman, G., Visintainer, R., Filosi, M., Riccadonna, S. & Furlanello, C. The HIM glocal metric and kernel for network comparison and classification. in 2015 IEEE International Conference on Data Science and Advanced Analytics (DSAA) 1–10 (IEEE, 2015).

36. Domingo-Gonzalez, R. et al. Diverse homeostatic and immunomodulatory roles of immune cells in the developing mouse lung at single cell resolution. Elife 9, e56890 (2020).

37. Zanini, F. et al. Progressive Increases in Mesenchymal Cell Diversity Modulate Lung Development and are Attenuated by Hyperoxia. bioRxiv (2021).

38. Zanini, F. et al. Phenotypic diversity and sensitivity to injury of the pulmonary endothelium during a period of rapid postnatal growth. bioRxiv (2021).

39. Schaum, N. et al. Single-cell transcriptomics of 20 mouse organs creates a Tabula Muris: The Tabula Muris Consortium. Nature 562, 367 (2018).

40. Stoeckius, M. et al. Simultaneous epitope and transcriptome measurement in single cells. Nature methods 14, 865–868 (2017).

41. Stuart, T., Srivastava, A., Lareau, C. & Satija, R.J.B. Multimodal single-cell chromatin analysis with Signac. BioRxiv (2020).

42. Wiener, N. Extrapolation, interpolation, and smoothing of stationary time series: with engineering applications, (MIT press Cambridge, MA, 1964).

43. Gentle, J.E.J.S.t.i.s., Springer, New York, NY, doi. Matrix algebra. Springer texts in statistics, Springer, New York, NY 10, 978–0 (2007).

44. Yin, C. & Yau, S.S.-T.J.J.o.c.b. A Fourier characteristic of coding sequences: origins and a non-Fourier approximation. Journal of computational biology 12, 1153–1165 (2005).

45. Bullard, J.H., Purdom, E., Hansen, K.D. & Dudoit, S. Evaluation of statistical methods for normalization and differential expression in mRNA-Seq experiments. BMC bioinformatics 11, 1–13 (2010).

46. Risso, D., Schwartz, K., Sherlock, G. & Dudoit, S. GC-content normalization for RNA-Seq data. BMC bioinformatics 12, 480 (2011).

47. Anders, S. & Huber, W. Differential expression analysis for sequence count data. Nature Precedings, 1–1 (2010).

## Reference

1. Chu, L.-F. et al. Single-cell RNA-seq reveals novel regulators of human embryonic stem cell differentiation to definitive endoderm. Genome biology 17, 1–20 (2016).

2. Zeisel, A. et al. Molecular architecture of the mouse nervous system. Cell 174, 999-1014. e22 (2018).

3. Schaum, N. et al. Single-cell transcriptomics of 20 mouse organs creates a Tabula Muris: The Tabula Muris Consortium. Nature 562, 367 (2018).

4. Baron, M. et al. A single-cell transcriptomic map of the human and mouse pancreas reveals inter-and intra-cell population structure. Cell systems 3, 346-360. e4 (2016).

5. Deng, Q., Ramsköld, D., Reinius, B. & Sandberg, R.J.S. Single-cell RNA-seq reveals dynamic, random monoallelic gene expression in mammalian cells. Science 343, 193–196 (2014).

6. Fan, X. et al. Single-cell RNA-seq transcriptome analysis of linear and circular RNAs in mouse preimplantation embryos. Genome biology 16, 1–17 (2015).

7. Goolam, M. et al. Heterogeneity in Oct4 and Sox2 targets biases cell fate in 4-cell mouse embryos. Cell 165, 61–74 (2016).

8. Klein, A.M. et al. Droplet barcoding for single-cell transcriptomics applied to embryonic stem cells. Cell 161, 1187–1201 (2015).

9. Li, H. et al. Reference component analysis of single-cell transcriptomes elucidates cellular heterogeneity in human colorectal tumors. Nature genetics 49, 708–718 (2017).

10. La Manno, G. et al. Molecular diversity of midbrain development in mouse, human, and stem cells. Cell 167, 566-580. e19 (2016).

11. Marques, S. et al. Oligodendrocyte heterogeneity in the mouse juvenile and adult central nervous system. Science 352, 1326–1329 (2016).

12. Pollen, A.A. et al. Low-coverage single-cell mRNA sequencing reveals cellular heterogeneity and activated signaling pathways in developing cerebral cortex. Nature biotechnology 32, 1053–1058 (2014).

13. Tasic, B. et al. Adult mouse cortical cell taxonomy revealed by single cell transcriptomics. Nature neuroscience 19, 335–346 (2016).

14. Yan, L. et al. Single-cell RNA-Seq profiling of human preimplantation embryos and embryonic stem cells. Nature structural molecular biology 20, 1131–1139 (2013).

15. Zheng, G.X. et al. Massively parallel digital transcriptional profiling of single cells. Nature communications 8, 1–12 (2017).

16. Domingo-Gonzalez, R. et al. Diverse homeostatic and immunomodulatory roles of immune cells in the developing mouse lung at single cell resolution. Elife 9, e56890 (2020).

17. Zanini, F. et al. Progressive Increases in Mesenchymal Cell Diversity Modulate Lung Development and are Attenuated by Hyperoxia. bioRxiv (2021).

18. Zanini, F. et al. Phenotypic diversity and sensitivity to injury of the pulmonary endothelium during a period of rapid postnatal growth. bioRxiv (2021).

19. Schmidt, M.N., Winther, O. & Hansen, L.K. Bayesian Non-negative Matrix Factorization. in Independent Component Analysis and Signal Separation, Vol. 5441 (eds. Adali, T., Jutten, C., Romano, J.M.T. & Barros, A.K.) 540–547 (Springer Berlin Heidelberg, Berlin, Heidelberg, 2009).

20. Spearman, C. “General Intelligence” Objectively Determined and Measured, 9 (Appleton-Century-Crofts, East Norwalk, CT, US, 1961).

21. Hyvärinen, A. & Oja, E. Independent component analysis: algorithms and applications. Neural Networks 13, 411–430 (2000).

22. Dasgupta, S. Experiments with Random Projection. 1301.3849 [cs, stat] (2013).

23. Ross, D.A., Lim, J., Lin, R.-S. & Yang, M.-H. Incremental Learning for Robust Visual Tracking. International Journal of Computer Vision 77, 125–141 (2008).

24. Tenenbaum, J.B., Silva, V.d. & Langford, J.C. A Global Geometric Framework for Nonlinear Dimensionality Reduction. Science 290, 2319–2323 (2000).

25. Szubert, B., Cole, J.E., Monaco, C. & Drozdov, I. Structure-preserving visualisation of high dimensional single-cell datasets. Scientific Reports 9, 8914 (2019).

26. Schölkopf, B., Smola, A. & Müller, K.-R. Kernel principal component analysis. (eds Gerstner, W., Germond, A., Hasler, M. & Nicoud, J.-D.) 583–588 (Springer, 1997).

27. Blei, D.M., Ng, A.Y. & Jordan, M.I. Latent Dirichlet Allocation. Journal of Machine Learning Research 3, 993–1022 (2003).

28. Lin, C.-J. Projected Gradient Methods for Nonnegative Matrix Factorization. Neural Computation 19, 2756–2779 (2007).

29. Roweis, S.T. & Saul, L.K. Nonlinear Dimensionality Reduction by Locally Linear Embedding. Science 290, 2323–2326 (2000).

30. Cichocki, A. & Phan, A.-H. Fast Local Algorithms for Large Scale Nonnegative Matrix and Tensor Factorizations. IEICE TRANSACTIONS on Fundamentals of Electronics, Communications and Computer Sciences E92-A, 708–721 (2009).

31. Lee, D.D. & Seung, H.S. Learning the parts of objects by non-negative matrix factorization. Nature 401, 788–791 (1999).

32. Pascual-Montano, A., Carazo, J.M., Kochi, K., Lehmann, D. & Pascual-Marqui, R.D. Nonsmooth nonnegative matrix factorization (nsNMF). IEEE Transactions on Pattern Analysis and Machine Intelligence 28, 403–415 (2006).

33. Moon, K.R. et al. Visualizing structure and transitions in high-dimensional biological data. Nature Biotechnology 37, 1482–1492 (2019).

34. Halko, N., Martinsson, P.G. & Tropp, J.A. Finding Structure with Randomness: Probabilistic Algorithms for Constructing Approximate Matrix Decompositions. SIAM Review 53, 217–288 (2011).

35. Laurberg, H., Christensen, M.G., Plumbley, M.D., Hansen, L.K. & Jensen, S.H. Theorems on Positive Data: On the Uniqueness of NMF. in Computational Intelligence and Neuroscience (2008).

36. Dueck, D., Frey, B.J., Dueck, D. & Frey, B.J. Probabilistic sparse matrix factorization. (2004).

37. Amodio, M. et al. Exploring single-cell data with deep multitasking neural networks. Nature Methods 16, 1139–1145 (2019).

38. Kim, H. & Park, H. Sparse non-negative matrix factorizations via alternating non-negativity-constrained least squares for microarray data analysis. Bioinformatics 23, 1495–1502 (2007).

39. Zou, H., Hastie, T. & Tibshirani, R. Sparse Principal Component Analysis. Journal of Computational and Graphical Statistics 15, 265–286 (2006).

40. Li, P., Hastie, T.J. & Church, K.W. Very sparse random projections. 287–296 (Association for Computing Machinery, 2006).

41. Belkin, M. & Niyogi, P. Laplacian Eigenmaps and Spectral Techniques for Embedding and Clustering. in Advances in Neural Information Processing Systems 14 (eds. Dietterich, T.G., Becker, S. & Ghahramani, Z.) 585–591 (MIT Press, 2002).

42. Maaten, L.v.d. & Hinton, G. Visualizing Data using t-SNE. Journal of Machine Learning Research 9, 2579–2605 (2008).

43. McInnes, L., Healy, J. & Melville, J. UMAP: Uniform Manifold Approximation and Projection for Dimension Reduction. 1802.03426 [cs, stat] (2018).

44. Wang, D. & Gu, J. VASC: dimension reduction and visualization of single cell RNA sequencing data by deep variational autoencoder. bioRxiv, 199315 (2017).

45. Chen, S., Hua, K., Cui, H. & Jiang, R. VPAC: Variational projection for accurate clustering of single-cell transcriptomic data. BMC Bioinformatics 20, 0 (2019).

46. Pierson, E. & Yau, C. ZIFA: Dimensionality reduction for zero-inflated single-cell gene expression analysis. Genome Biology 16, 241 (2015).

47. Saelens, W., Cannoodt, R., Todorov, H. & Saeys, Y.J. A comparison of single-cell trajectory inference methods. Nature biotechnology 37, 547–554 (2019).

